# Collective dynamical regimes predict invasion success and impacts in microbial communities

**DOI:** 10.1101/2024.02.05.579032

**Authors:** Jiliang Hu, Matthieu Barbier, Guy Bunin, Jeff Gore

## Abstract

Invasions of microbial communities by species such as pathogens can have significant impacts on ecosystem services and human health^1–9^. Predicting the outcomes of these invasions, however, remains a challenge. Various theories propose that these outcomes depend on either characteristics of the invading species^10–12^ or attributes of the resident community^13–16^, including its composition and biodiversity^3,17–19^. Here we used a combination of experiments and theory to show that the interplay between dynamics, interaction strength, and diversity determine the invasion outcome in microbial communities. We found that the communities with fluctuations in species abundance are both more invasible and more diverse than stable communities, leading to a positive diversity-invasibility relationship among communities assembled in the same environment. As predicted by theory, increasing interspecies interaction strength and species pool size leads to a decrease of invasion probability in our experiment. Although diversity-invasibility relationships are qualitatively different depending upon how the diversity is changed, we provide a unified perspective on the diversity-invasibility debate by showing a universal positive correspondence between invasibility and survival fraction of resident species across all conditions. Communities composed of strongly interacting species can exhibit an emergent priority effect in which invader species are less likely to colonize than species in the original pool. However, in this regime of strong interspecies interactions, if an invasion is successful, it causes larger ecological effects on the resident community than when interactions are weak. Our results demonstrate that the invasibility and invasion effect are emergent properties of interacting species, which can be predicted by simple community-level features.

## Introduction

Ecological invasions, characterized by the spread of non-native species into novel environments, have significant consequences for biodiversity, ecosystem function, and habitat resilience ^20^. Over decades, ecologists have sought to unravel the myriad factors influencing why some species invade successfully, and why some of those have large impacts on resident species communities, while others do not. Ecologists have posited a range of determinants, from the fitness and adaptability of the invaders to the resilience and composition of native communities ^21–23^. Among studies focusing on the invader species, many have sought to identify traits, such as growth and dispersal strategies, that may shape invasion outcomes^10^. Others have emphasized the role of the invaders’ initial population size in the likelihood of establishment and spread^11,12^. Yet others have emphasized interactions with resident species, e.g. the enemy release hypothesis suggesting that invasive species often succeed in new environments because they lack consumers or pathogens^16^. This has led to research on how properties of resident communities as a whole can determine the invasion outcome. For instance, the biotic resistance hypothesis suggests that communities with high native biodiversity are more resistant to invasion than less diverse communities, due to more efficient resource use or presence of natural enemies, but it is not consistently supported by empirical results^13–15,24^. Beyond the characteristics of invader species and resident communities, environmental conditions have been shown to play a crucial role in shaping the invasion outcome^20^. Theories such as the storage effect and the fluctuating resource availability hypothesis posit that environmental disturbances and fluctuations might favor invader species in specific periods^25–27^.

More recently, the issue of ecological invasion has become salient in the study of microbial communities, ranging from soil and aquatic ecosystems to the human body^1–7^. These invasions can have profound impacts on ecosystem services and human health^1,2,4,5^. Pathogenic microbes can invade host-associated microbial communities, leading to infections and diseases^4,8,9^. For example, the invasion of pathogenic microbes *C. difficile* into the gut microbiota can lead to severe diseases including diarrhea and colitis^8,28^. Understanding the mechanisms underlying invasion success and ecological consequences can help inform strategies for disease prevention, as well as the development of targeted therapies to control invasive pathogens^28,29^. Similar to larger-scale ecological systems, it has been suggested that microbial communities with higher diversity are less likely to be invaded because diverse resident species may occupy all available niches and resources, leaving less room for invaders^3,17–19^. Furthermore, it was shown that facilitative and competitive interactions between microbes can favor and prevent successful invasions, respectively^17,30–32^. Parallel to observations in macro-organisms, external disruptions such as antibiotic interventions or nutrient level shifts can heighten the vulnerability of microbial communities to invasions^1,33–35^.

While research in microbial invasions has made significant strides, it remains unclear what characteristics of resident communities determine the success and impacts of an invasion^2,3,36,37^. Species diversity is an easily measured indicator, but its relationship to invasibility may not be straightforward, whereas species interactions are likely important but often difficult to quantify. A more rarely emphasized property is the residents’ dynamics: are they at equilibrium or fluctuating? It is not obvious that we can characterize dynamics at the level of the community; yet, building upon the groundbreaking work of Robert May, ecologists have explored the possibility of community-wide emergent dynamics, that can be classified into only a few qualitatively distinct regimes. and predicted from a few macroscopic parameters^14,38–43^. In a recent study^40^, we experimentally assembled communities from various pools of microbial species in different conditions, and confirmed that simple community-level features, including species pool size and interaction strength, determined distinct dynamical regimes, characterized by the fraction of surviving species and the emergence of persistent abundance fluctuations over time.

Here, we perform invasion experiments in diverse assembled microbial communities, and first observe that the foremost predictor of invasion outcomes appears to be the dynamical state of the resident community. We then use a combination of experiments and theory, exploring several dynamical regimes and spanning their control parameters (species pool size and interaction strength) to show that, taken together, they explain many features of invasibility and invasion effects. Communities of weakly interacting species reach a stable composition, where a fraction of the initial species pool survives, and further invasions display the same fraction of successes, only weakly perturbing resident species. Richer species pools and stronger interactions can give rise to fluctuating states, where species abundances fluctuate over time. We found that these fluctuating communities are more invasible and diverse than stable communities, leading to a positive diversity-invasibility relationship among communities assembled in the same environment. Finally, communities with strong interactions may also reach stable states with lower diversity, whose experimental phenomenology evokes the presence of alternative stable states: invasions succeed more rarely than predicted by survival fraction, but strongly impact the resident community when they do. The lower probability of invasion suggests a priority effect, whereby earlier invaders preclude later ones from growing from small abundances, leading to situations where the sequence and timing of species introduction can influence invasion success^14,44,45^.

Studying invasions through the prism of community-wide dynamical regimes allows us to connect several strands of ecological thinking, regarding what counts as a successful invasion, when factors such as population size and history matter, and what consequences invasions have on resident community structure and functioning^46,47^. Furthermore, it helps clarify the hypothesis that increased community diversity results in reduced invasion probability due to fewer available niches^3,17–19^. Within fixed conditions (given the same initial species pool size and environment), more diverse communities tend to be found in fluctuating states, and are actually more likely to be invaded. Depending on how we change conditions, e.g. increasing species pool or reducing interaction strength, diversity may positively or negatively correlate with invasibility, providing one explanation for inconsistent observations^48–50^. Throughout, the fraction of surviving species remains a better predictor, displaying a universal positive correspondence with invasibility across all conditions, modulated by the presence of priority effects. Our results demonstrate that both invasibility and invasion effects are emergent properties, shaped by the interactions of resident species, which can be predicted by simple community-level features.

## Results and discussion

To experimentally characterize invasions in microbial communities, we built 17 different synthetic communities of size *S* = 20 using a library of 80 bacterial isolates from river and terrestrial environments (Fig. 1a and Supplementary Fig. 1). We exposed each community to daily cycles of growth and dilution into fresh media, with dispersal from the species pool (*S* = 20) to mimic species dispersal in natural habitats (Fig. 1a). After six days of culturing, each community was exposed to an invader species (Fig. 1a) and we continued to culture the communities for another 6 days with dispersal of all species on each dilution cycle (Fig. 1a, b). For each resident community, we performed 7-9 independent invasion tests with different invader species on day 6, and monitored the growth of the invader and resident species (Fig. 1b). Analyzing species abundances through 16S sequencing, we found that 7% ± 2% of invasion tests were successful (relative invader abundance exceed extinction threshold 8×10^-4^ on the last day 12) (Fig. 1c and Supplementary Fig. 2). Although diverse ecosystems are typically thought to be more resistant to invaders^3,17–19^, our experimental results display a significant (p=0.036) positive correlation between invasion probability and community diversity (correlation coefficient=0.51, Fig. 1c). Among communities of low diversity (2 – 5 surviving species) only 3% ± 2% of invasions were successful, whereas among communities of high diversity (6 – 9 surviving species) 13% ± 5% of invasions were successful. We therefore find that less diverse communities may resist invasions better than highly diverse ones under the same initial species pool size and nutrient conditions.

**Fig 1.**
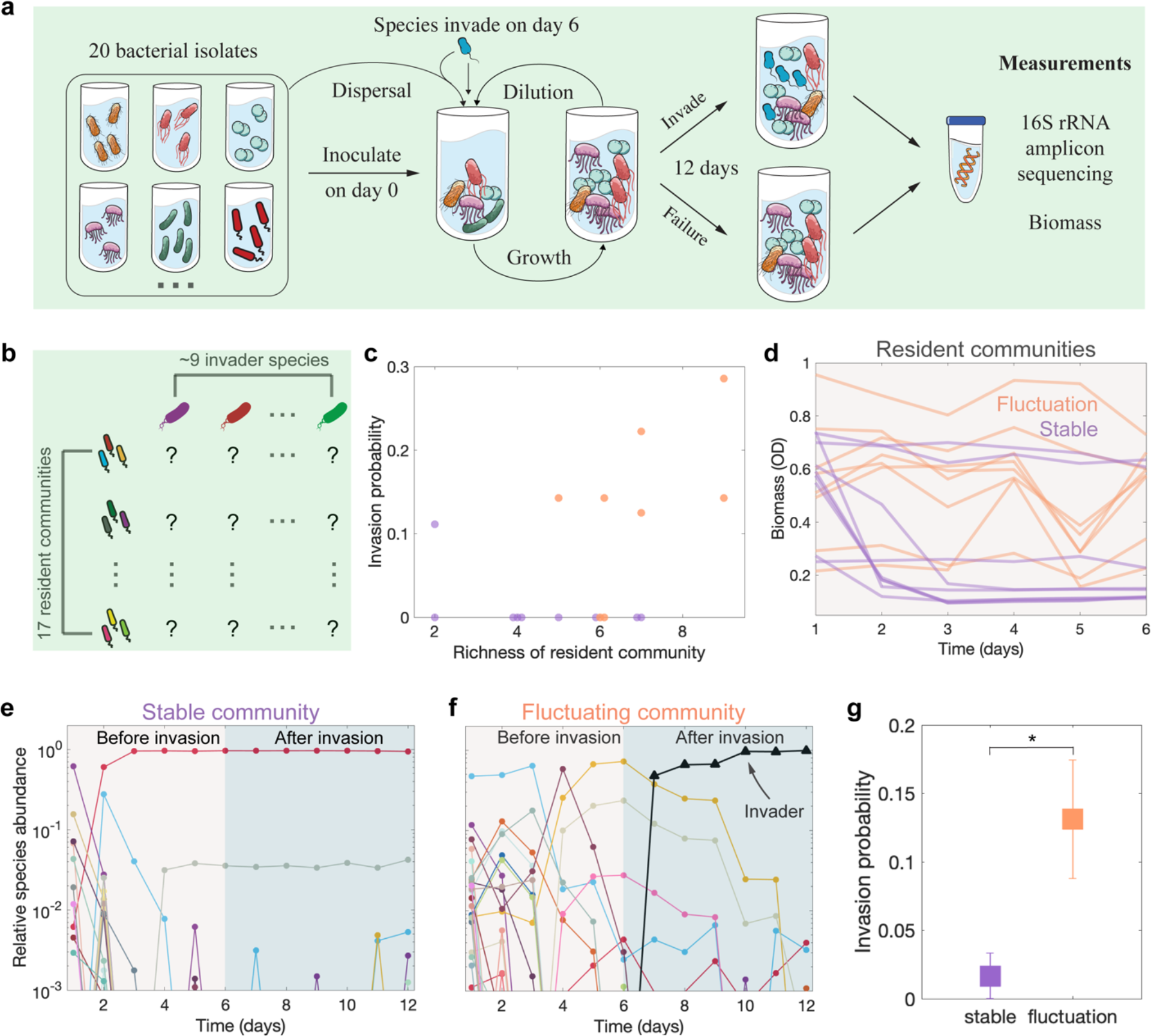
Experiments using synthetic microbial communities show that the invasion probability in fluctuating communities is higher than stable ones, leading to a positive diversity-invasibility relationship. **a**, We used a library of bacteria to generate different synthetic communities with *S*=20 species in the pool (under “high” nutrient conditions; see supplement).

To better understand why the more diverse communities were more invasible, we next quantified the dynamics of the resident communities before invasion. We found that just under half (8 / 17) of the resident communities displayed persistent fluctuations in biomass and species composition, with the remainder reaching stable community states (Fig. 1d-f and Supplementary Fig. 3-12). We found that biomass fluctuations were highly correlated with species abundance fluctuations (Supplementary Fig. 12) and the classification of stable and fluctuating communities was robust to different methods (Supplementary Fig. 12). Consistent with our previous results, we found that the diversity of fluctuating communities is approximately twice the diversity in stable communities (Fig. 1c) ^40^. Given this higher diversity in fluctuating communities, we next analyzed the invasibility of communities separately for the stable and fluctuating communities to determine if this could be driving the positive diversity-invasibility relationship that we observed. Indeed, we detected 8 successful invasions out of 61 invasion tests to fluctuating communities, while there was only one single successful invasion out of 60 invasion tests to stable communities (Supplementary Fig. 2). Our results therefore show that the probability to successfully invade fluctuating communities (13% ± 4%) is statistically ∼8 fold larger than the probability of invading stable communities (1.7% ± 1.7%) (Fig. 1g). Our experimental tests of invasion demonstrate that, for fixed species pool size and conditions (including species interaction strength), more diverse communities are more invasible because fluctuating communities are both more diverse and more susceptible to invasion.

To gain insight into these surprising relationships between diversity, stability, and invasibility, we next studied invasions in the well-known generalized Lotka-Volterra (gLV) model, modified to include dispersal from a species pool:

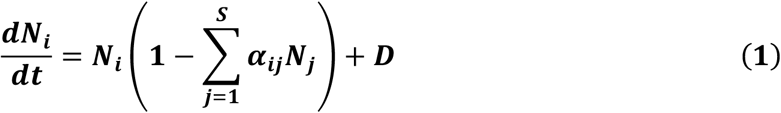

where *Ni* is the abundance of species *i* (normalized to its carrying capacity), *αij* is the interaction strength that captures how strongly species *j* inhibits species *i* (with self-regulation *αii* = 1), and *D* is the dispersal rate. We simulated the dynamics of communities with different species pool sizes *S* and interaction matrices. We sampled the interaction strength from a uniform distribution *U* [*0, 2*<*αij*>], where <*αij*> is the mean interaction strength between species (predictions of this model are insensitive to the particular distribution chosen^40^). Modeling species interactions as a random interaction network captures species heterogeneity without assuming any particular community structure ^14,38–40^. We introduced invaders into resident communities at *t*=10^3^ and continued to simulate the dynamics until *t*=2×10^3^ to determine to invasion outcome.

Our simulations revealed rich dynamics and invasion outcomes under strong interaction strength between species (Fig. 2a). Some successful invasions cause dramatic effect on the structures of resident communities, whereas other invasions only yield weak change in communities (Fig. 2a). Consistent with our experimental results (Fig. 1c and 1g), we found a positive correlation between invasion probability and richness (Fig. 2b), which is because fluctuating communities exhibit larger invasion probability than stable communities under the same conditions (Fig. 2c). Our simulation results with the Lotka-Volterra model also predict that the invasion probability decreases when interaction strength <*αij*> and the species pool size *S* increase (Fig. 2d-f). It is important to note that although fluctuating communities exhibit smaller invasion resistance than stable communities under the same conditions, stable communities can still yield smaller invasion resistance under weaker interaction strength <*αij*> or smaller species pool size *S* (Fig. 2d-f). The Lotka-Volterra model therefore explains why our diverse and fluctuating communities are susceptible to species invasion and makes new predictions regarding how invasibility would change with the size of the species pool and the strength of interspecies interactions (Fig. 2d-f).

**Fig 2.**
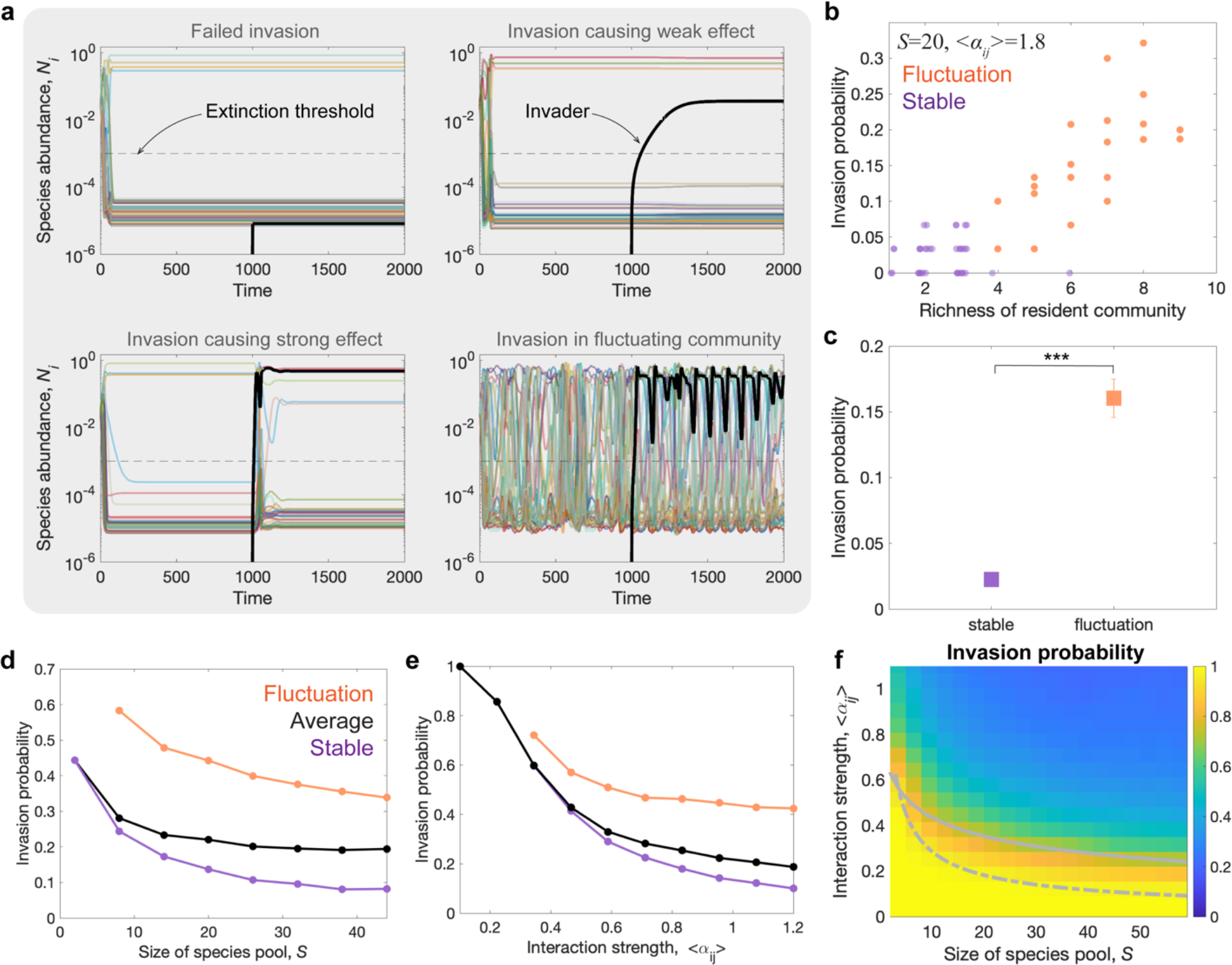
The Lotka-Volterra model predicts a decrease of invasion probability when stability, interaction strength and species pool size of resident communities increase. **a**, Representative time series of species abundance in simulation show diverse invasion dynamics and outcome: invader species failed to grow in the community (top left panel, the black curve represents invader); invader grow and only cause small effect on community composition (top right panel); invader successfully invade and cause large change on community composition (bottom left panel); invasion to a fluctuating resident community (bottom right panel) (<*αij*>=0.6, *S*=32). **b**, Consistent with experiments (Fig. 1c), the invasion probability of simulated resident communities positively correlates with their richness, which arises because fluctuating communities are more diverse and more invasible. **c**, The invasion probability to fluctuating resident communities is statistically higher than that to stable communities (p < 0.001). **d**, Increasing species pool size leads to a decrease in invasion probability. Fluctuating communities (orange points) exhibit higher invasion probability than stable communities (purple points). **e**, Increasing interaction strength leads to a decrease in invasion probability. **f**, Increasing species pool size and interaction strength leads to a decrease in invasion probability. The curves and color maps depict the mean value over 1000 simulations.

To experimentally test the predicted dependence of invasion resistance on interaction strength and species pool size, we tuned the inter-species interaction strength by tuning the concentration of supplemented glucose and urea in the culture medium ^40,51,52^. As discussed in our previous work ^40,51,52^, increasing the concentration of supplemented glucose and urea leads to stronger strength of competitive interactions between bacterial species due to extensive modification of the media (e.g. pH). We measured the invasion of ∼9 invader species to 15 synthetic resident communities under low nutrient (weak interaction) and 25 communities under high nutrient (strong interaction) conditions. Consistent with our theoretical predictions, we found that increasing interaction strength leads to an increase of invasion resistance in resident communities (Fig. 3a). Specifically, the invasion probability was 7% ± 2% in high nutrient conditions (strong interaction), 8 fold lower than the invasion probability of 56 % ± 8% observed in low nutrient conditions (weak interaction) (Fig. 3a). We also decreased the species pool size from *S*=20 to *S*=12 and found that invasion probability increased to 85 % ± 6 % from 56 % ± 8% in low nutrient conditions (Fig. 3b), consistent with our theoretical predictions. Our theory and experiment both indicate that increasing either interaction strength or species pool size leads to a decrease in community invasibility ^3,14,17–19^.

**Fig 3.**
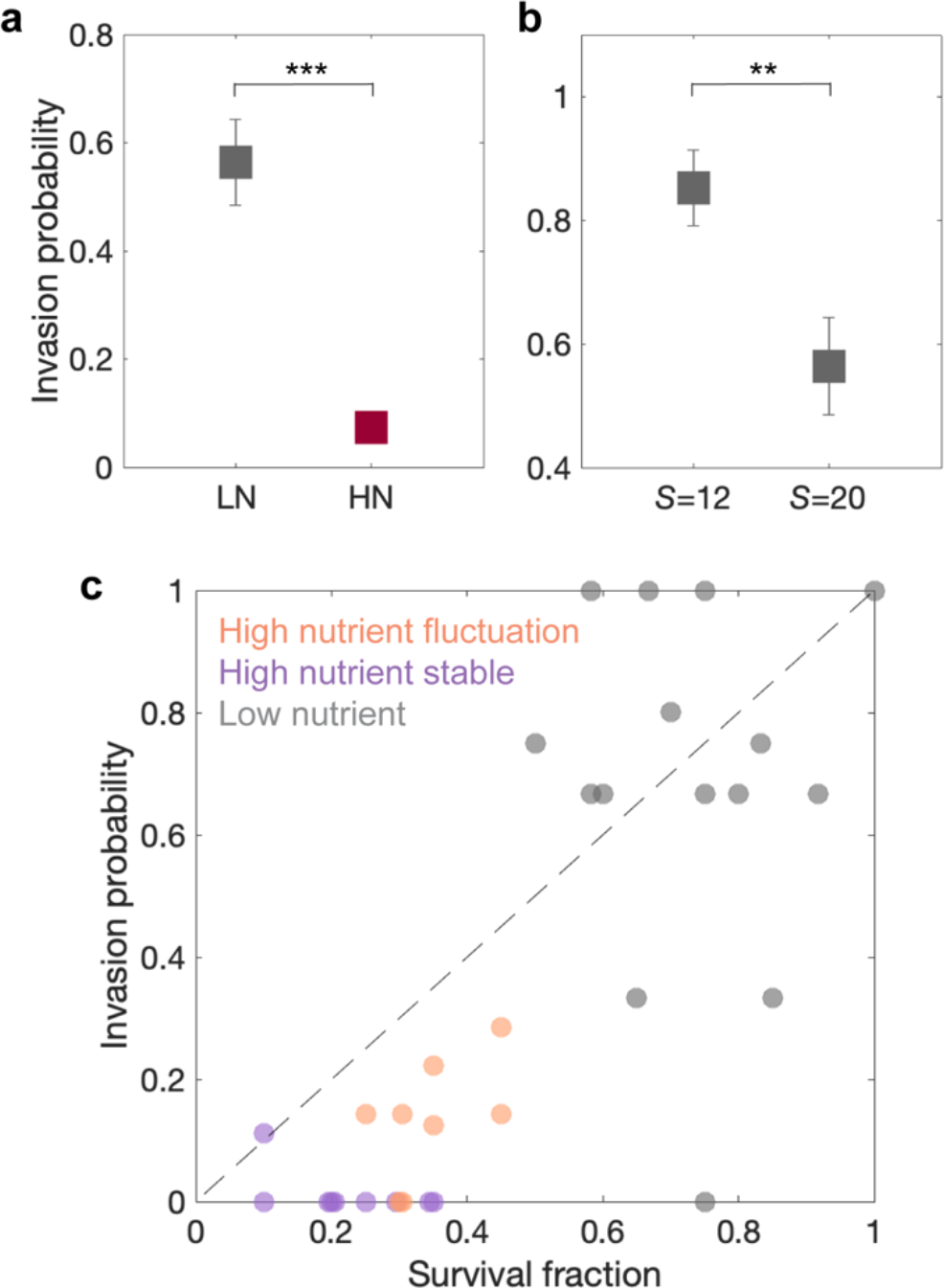
Increasing interaction strength and species pool size leads to higher invasion resistance of resident communities in experiment. **a**, The invasions to resident communities under low nutrient (weak interaction) exhibit statistically higher invasion probability than communities under high nutrient (strong interaction) (p < 0.001, the number of invasion tests is n=120 (39) for high (low) nutrient). **b**, The invasions to resident communities under smaller species pool size (*S*=12) exhibit statistically higher invasion probability than communities under larger species pool size (*S*=20) (p = 0.007, the number of invasion tests is n=39 (34) for *S*=20 (12), all communities were cultured under low nutrient). Error bars represent s.e.m.. **c**, The invasion probability positively correlates with survival fraction (before invasion) across different communities and nutrient conditions (each point represents one community; correlation coefficient is 0.82). The points corresponding to communities under high nutrient are below the diagonal line, showing the invasion probability of communities under high nutrient are generally smaller than their survival fraction, which indicates the priority effect under strong interaction strength.

To unify different invasibility-richness relationships in the experiments depending upon how the richness is changed (by varying interaction strength, species pool size, or dynamical regime) (Supplementary Fig. 13), we next analyzed the dependence of invasion probability on the survival fraction of species in resident communities, defined as the fraction of species in the initial pool which survive the assembly process (on day 6 before invasion). The results show a strongly positive correlation of invasibility on survival fraction, where the correlation coefficient is 0.82 (Fig. 3c). Microbial communities cultured in low nutrient (weak interaction) media display both a larger invasion probability and larger survival fraction than communities under high nutrient (strong interaction) (Fig. 3c). Furthermore, fluctuating communities which are easier to be successfully invaded also exhibit larger survival fraction than stable communities under the same conditions (Fig. 3c and 1c). These results demonstrate that the survival fraction can serve as a unifying predictor of the invasibility of a resident community. Although it has been suggested that microbial communities with higher diversity are less likely to be invaded because they leave less available niches and resources for invaders^3,17–19^, our results indicate that this is only true when the diversity is increased by increasing the size of the species pool (Fig. 3c and 1c). However, if diversity is modulated by a change in interaction strength or stability then more diverse communities are instead more invasible.

Despite our experimentally observed correspondence between invasion probability and survival fraction, we noted that the invasion probability for communities under high nutrient are usually lower than their survival fraction (majority of points on the bottom left (high nutrient) are below the diagonal line on Fig. 3c). In ecology, the priority effect refers to the phenomenon in which the community structure is influenced by the order and timing of species’ arrival in a community^44,45^. We interpret the mismatch between invasion probability and survival fraction under high nutrient as evidence of priority effects in the community assembly under strong interactions in our experiment. Under weak interactions, the colonization probability of invader species is similar with the probability of a species in the initial pool surviving the process of community assembly, whereas invader species are statistically less likely to survive in the communities than the species in the initial pool under strong interactions (Fig. 3c)^14^. The emergent priority effect in communities composed of strongly interacting species could be explained by alternative stable states tending to inhibit the growth of invaders at low abundance^14,40^.

To understand the reason for different diversity-invasibility relationships when varying interaction strength, species pool size or dynamical regime (Fig. 1c, Fig. 2 and Fig. 3), we sampled resident communities along different paths on the phase diagram (Fig. 2f). We simulated invasions to these resident communities and found different diversity-invasibility relationship along different paths (Fig. 4a). The results show a positive diversity-invasibility relationship when only varying interaction strength while fixing species pool size, or randomly sampling communities under the same parameters of species pool size and interactions (Fig. 4a). On the contrary, a reversed negative or non-monotonic diversity-invasibility relationship was observed when varying species pool size while fixing interaction strength (Fig. 4a). Despite these conflicting diversity-invasibility relationships, after scaling richness with species pool size to get the survival fraction, we found that all communities collapsed to a universal line in which the invasion probability is approximately equal to the survival fraction (Fig. 4b). The deviation from the exact collapse in small survival fraction regime (bottom left of Fig. 4b) indicates priority effect under strong interaction. Our results indicate that survival fraction determines invasibility, whereas diversity-invasibility relationship can be qualitatively different depending upon the origin of different diversity in different communities.

**Fig 4.**
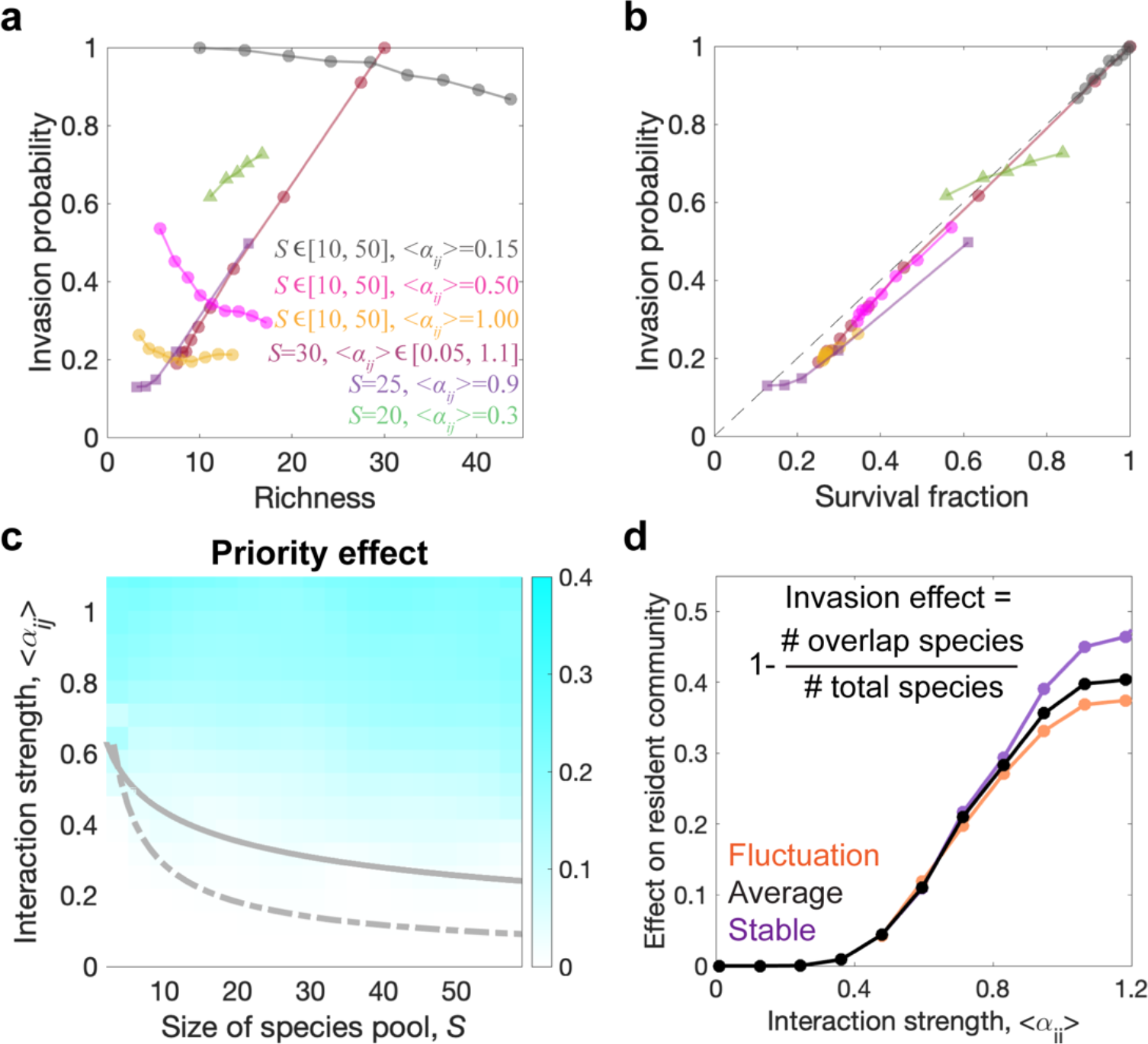
The Lotka-Volterra model predicts a universal correspondence between invasion probability and survival fraction, the emergence of priority effects and stronger invasion effects when increasing interaction strength. **a**, The dependence of invasion probability on final richness of resident communities is qualitatively different depending upon how the richness is changed. Invasion probability positively correlates with richness when varying interaction strength or when randomly sampling communities with a fixed species pool size and interaction strength distribution. Invasion probability can decrease with community diversity when varying species pool size. **b**, Invasion probability is approximately equal to the survival fraction of species in the resident communities, no matter how we change richness, species pool or interaction strength. **c**, Increasing species pool size and interaction strength leads to the emergence of priority effect, where the invasion probability of resident communities is smaller than their species survival fraction. **d**, Successful invasions cause larger effect on species composition in the resident communities under stronger interaction strength. The curves depict the mean value over 1000 simulations.

The emergence of the priority effect in experiments (Fig. 3c) was also found in the Lotka-Volterra model under different regimes of interaction strength and species pool size. We quantified the priority effect by calculating the difference between survival fraction of resident species and the invasion probability of species that invade after the resident communities have assembled, where the difference was normalized by survival fraction (Fig. 4c). We found there is no clear priority effect in the small species pool size and weak interaction regime, where species in the initial pool and invader species display similar probability of colonizing in the communities (Fig. 4c). Consistent with our experimental results, increasing species pool size and interaction strength in the model leads to the emergence of priority effect in the phase where communities reach fluctuation or alternative stable states (Fig, 4c). We found the priority effect originated from alternative stable states or limit cycle oscillations in the strongly interacting phase, whereas chaotic fluctuations display no significant priority effect in simulation (Supplementary Fig. 14), which can be explained by its ergodicity^41,43^.

We also investigated the idea that successful invasions can cause strong or weak effects on resident community structure depending on how invaders interact with resident species^14,30,31^. Our simulations show that invasions cause stronger change on structure of resident communities under stronger interactions, where the invasion effect is measured as the proportion of change in surviving species before and after the invasion, calculated as 1 minus the ratio of overlapping species to total species (1 -(number of overlapping species / total number of species)). (Fig. 4d). To understand the effect of a successful invasion in the experiment, we analyzed the change of biomass and species composition before and after the invasions (Fig. 5). The community biomass displays relatively small changes after invasion under weak interactions (low nutrient regime, inset of Fig. 5a and Supplementary Fig. 6-7). In the strong interaction regime (high nutrient), we found that stable communities typically transitioned from low biomass states to high biomass states after successful invasions, whereas the biomass of fluctuating communities continued to fluctuate over a similar range (Fig. 5a-c and Supplementary Fig. 3-5). Averaging across both stable and fluctuating communities, we found that community biomass under strong interaction displayed a larger fold change (2.9 ± 0.8) after successful invasion than those under weak interaction (1.15 ± 0.03) (Fig. 5c). We defined the invasion effect as the fraction of surviving species that change before and after invasion. We compared the change of relative species abundance between invaded communities and control communities without adding invaders (Supplementary Fig. 15-18). This analysis on species abundances through 16S sequencing further indicated that successful invasions cause stronger change in the species compositions under strong interaction (53 % ± 6%) than weak interaction (39% ± 2%) (Fig. 5d), which is consistent with the simulation results with Lotka-Volterra model (Fig. 4d). The growth of invader species influences the community structure more dramatically when it has stronger interaction with other resident species, and the strong interplay between resident species can also cause stronger secondary effect on other resident species when their abundance change^14,47^.

**Fig 5.**
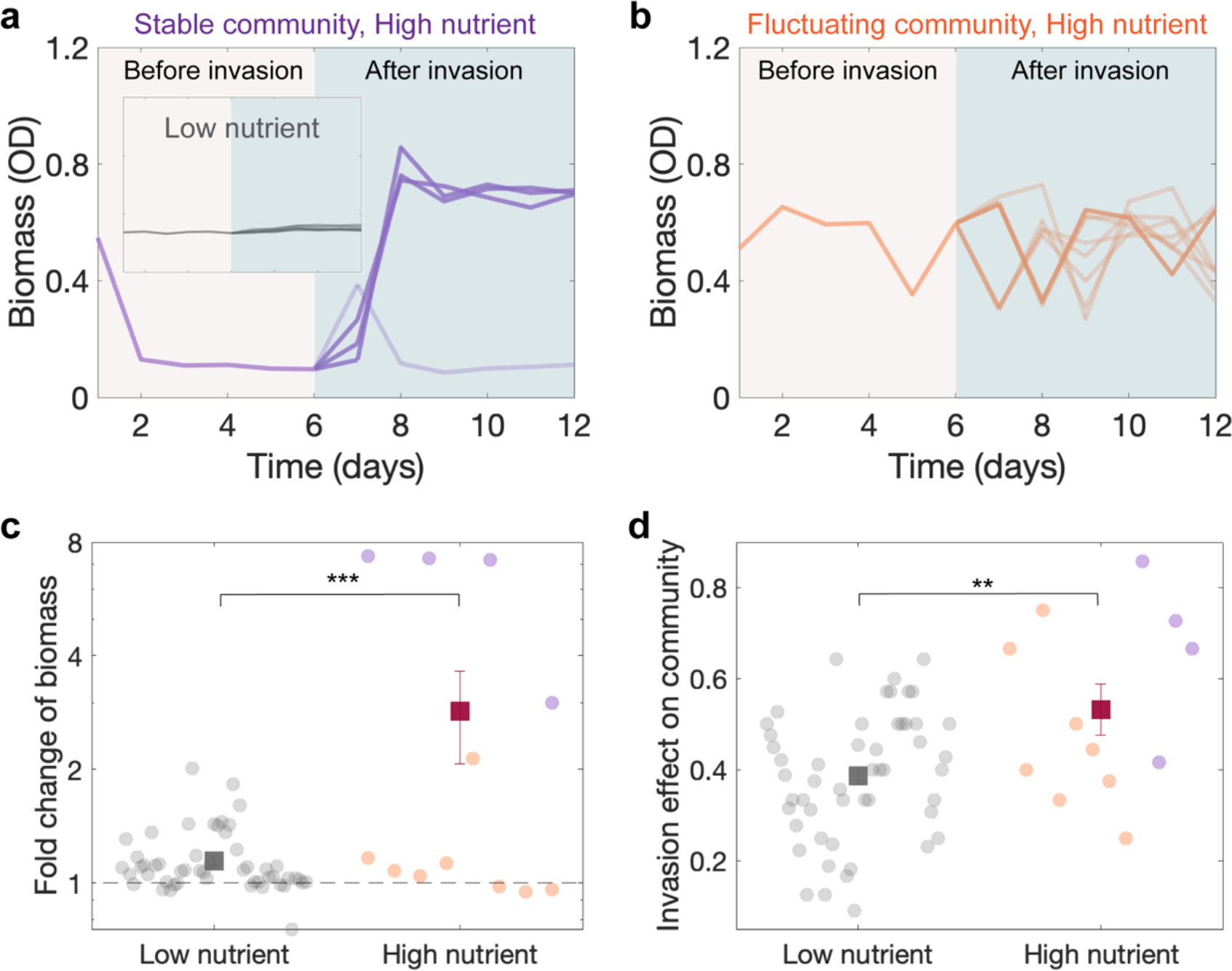
Increasing interaction strength leads to a stronger effect on resident communities under invasion success. **a**, The stable communities under high nutrient experienced a large increase in biomass after successful invasions (dark purple curves). Inset shows the invasions under low nutrient only cause weak effect on community biomass as compared with high nutrient. **b**, The time course of fluctuating community biomass under high nutrient before invasion and after invasion, where dark and light orange curves represent successful and failed invasions, respectively. **c**, The invasions to resident communities under low nutrient (weak interaction) cause statistically lower fold change of biomass than communities under high nutrient (strong interaction) (p < 0.001, the number of successful invasions is n=51 (11) for low (high) nutrient). The successful invasions statistically tend to increase the biomass of resident communities under different conditions. **d**, The invasions to resident communities under low nutrient (weak interaction) cause statistically lower effect on species composition change than communities under high nutrient (strong interaction) (p = 0.0038, the number of invasion tests is n=51 (11) for low (high) nutrient). Error bars represent s.e.m..

Although our study was primarily focused on community-level properties that determine invasibility and invasion effect, we also analyzed properties of the invader species that correlated with invasibility and invasion effect. Perhaps surprisingly, we did not observe a significant correlation between a species’ ability to invade and that species growth in monoculture (Supplementary Fig. 19). For example, Pseudomonas (invader 4) and Enterobacterales (invader 7) were the two most successful invader species (16/35 and 6/11 invasions, respectively), yet displayed growth in monoculture that was typical of the group of nine invaders that were tested. Bacillus (invader 6) with the highest growth rate among all invaders under both high and low nutrients, only display a small fraction of invasion success (2/37). Moreover, we also observed that Pseudomonas (invader 2) and Pedobacter (invader 3) could occasionally invade communities despite being subject to a strong Allee effect that prevented the species from growing from an initially small inoculum (Supplementary Fig. 17). Furthermore, we did not observe significant correlation between the invasion effect and invader properties either (Supplementary Fig. 20). Whether for the invasion probability of resident communities or different invaders, we found an absence of correlation between invasion probability and invasion effect (Supplementary Fig. 21). Taking these together, we therefore found that monoculture growth properties were surprisingly ineffective at predicting the success of a species as an invader, while resident community properties are the primary factors determining the invasion outcome.

Our findings show that invasibility and invasion effects can be statistically predicted by simple community-level features including the community’s dynamical regime, species pool size, and interaction strength. As predicted by our theory, increasing community diversity leads to stronger resistance to invaders only when varying species pool size and fixing community stability and environmental conditions (including interaction strength), which is consistent with the biotic resistance hypothesis^13–15^. We demonstrated that, when diversity is tied to increased dynamic fluctuations or reduced interaction strength, more diverse communities might instead exhibit decreased resistance to invasion (Fig. 1c and 3c). Our results emphasize that only by concurrently considering the effects of interaction strength and stability can the diversity of native communities be used to predict invasion resistance; diversity alone is insufficient for such predictions. By normalizing richness with species pool size, we obtained the survival fraction, a unified predictor that closely approximates invasion probability across different conditions (Fig. 3c and Fig. 4b). This survival fraction is influenced by factors such as species pool size, interaction strength, and stability (Fig. 2b, cand 3c).

Applying the insights developed here to natural communities requires that we draw a parallel between the three recognized types of diversity in ecology—alpha, beta, and gamma diversity— and the three species number variables we’ve investigated in our study: richness, survival fraction, and species pool size^53^. Specifically, richness and species pool size can be seen as analogs for alpha (local diversity) and gamma (regional diversity) diversities, respectively. Beta diversity, defined as the ratio between regional and local diversity, is the reciprocal of the survival fraction. Consequently, our discovery of a universal positive relationship between invasibility and survival fraction suggests an overarching negative correlation between invasibility and beta diversity. While directly measuring the survival fraction in natural communities can be challenging, the ratio of local richness to regional richness in natural habitats may serve as a reliable approximation of survival fraction^13,24,48,53^, acting as a singular predictor for invasion probability. That prediction is nevertheless affected by the presence or absence of priority effects. Building upon our earlier discoveries regarding emergent phases in communities^40^, our current work suggests that priority effects are most pronounced in the theoretically-predicted phase of alternative stable states, matching our empirical observations of stable states found under conditions of strong interactions and a large species pool (Fig. 2e and 3c).

Our invasion experiments in synthetic microbial communities under controlled conditions have shown that, prior to any other feature of invader or resident species, the qualitative dynamical regime of the resident community is a central factor that informs all other predictions. The distinct regimes that we found, and the relationships between various predictions, were all compatible with a theory governed only by a few community-level parameters of (pool) diversity and interaction strength. Future work is necessary to determine whether these community-level features can predict invasion outcomes across spatiotemporal scales, environmental conditions, and organism types.

Communities underwent serial daily dilutions with additional dispersal from the pool. We introduced invader species to the resident communities on day 6 and continued to apply daily dispersal of invaders. Community composition and total biomass were monitored via 16S sequencing and optical density (OD). **b**, We formed 17 resident communities with different sets of species (*S*=20). We added invader species outside the pool into the resident communities on day 6, and then measured the community compositions and biomass on day 12 to determine the outcome and effect of the invasions. **c**, The invasion probability of resident communities positively correlate with their richness (correlation coefficient=0.51). **d**, 8 out of the 17 resident communities reach fluctuation in biomass (orange) and the rest 9 communities reach stable states (purple). **e**, Representative time course of relative species abundance via 16S sequencing show the stable community was not invaded. **f**, Representative time course of relative species abundance shows the invader successfully invade and grow in the fluctuating community. **g**, The invasion probability to fluctuating resident communities is statistically higher than that to stable communities (p=0.016, the number of invasion tests is n=61 (60) for fluctuating (stable) communities. Error bars represent s.e.m..

