## Supplementary Materials for "Collective dynamical regimes predict invasion success and impacts in microbial communities"

**This PDF file includes:**

### Materials and Methods

Supplementary Figs. 1 to 21

### Materials and Methods

#### Bacterial isolates, media and culturing conditions

We constructed the library of 80 bacterial species from soil, tree leaves, and Charles River water samples taken near the campus of Massachusetts Institute of Technology (Supplementary Fig. 1). This library is phylogenetically diverse, with isolates coming from 24 different families among 5 phylums: Proteobacteria, Firmicutes, Bacteroidota, Actinobacteriota, and Cyanobacteria (Supplementary Fig. 1).

In the case of low interaction strength (low nutrients concentration) conditions, experimental communities were cultured in Base Medium (BM): $1\text{ gL}^{-1}$ yeast extract and $1\text{ gL}^{-1}$ soytone from Becton Dickinson, 10 mM sodium phosphate, 0.1 mM $\text{CaCl}_2$, 2 mM $\text{MgCl}_2$, $4\text{ mgL}^{-1}$ $\text{NiSO}_4$ and $50\text{ mgL}^{-1}$ $\text{MnCl}_2$, pH adjusted to 6.5. For high interaction strength (high nutrients concentration) conditions, we used BM supplemented with $5\text{ gL}^{-1}$ glucose and $4\text{ gL}^{-1}$ urea. All media were filter sterilized using Bottle Top Filtration Units (VWR). All of the chemicals were purchased from Sigma–Aldrich unless otherwise stated.

Both monocultures and communities of the bacterial isolates were grown in 96-deepwell plates (Deepwell plate 96/500$\mu\text{L}$; Eppendorf) covered with AeraSeal adhesive sealing films (Excel Scientific). The incubation temperature was 30 °C for all communities. The deepwell plates were shaken at 1,200 r.p.m. on Titramax shakers (Heidolph). To minimize evaporation, the plates were incubated inside custom-built acrylic boxes.

#### Pre-cultures, daily dilutions, dispersal, invasion, and biomass measurements

Before each experiment, pre-cultures were initiated by thawing the bacteria and inoculating individual species into 300 $\mu\text{L}$ of BM. The resulting monocultures were exposed to 5 daily cycles of growth and (30-fold) dilution into fresh media. At the beginning of each experiment, aliquots of these monocultures were mixed in equal volume proportions to form the synthetic communities. During the experiment, the monocultures were exposed to further dilution cycles and used to apply the daily dispersal into the synthetic communities as described below.

We created 40 different synthetic communities using randomly generated subsets of the library of isolates, each subset constituting the species pool (of size $S$) for each community. After mixing monocultures in equal volumes, each experimental community was initiated by inoculating 10 $\mu\text{L}$ of its initial mix of isolates into 300 $\mu\text{L}$ of BM. The resulting synthetic communities were cultured under serial dilution cycles with dispersal as follows. To apply a $10^{-5}$ dispersal rate, every 24hr monoculture aliquots of the species in each community pool were mixed at equal volumes, and then diluted by a $10^3$ factor before inoculating 6$\mu\text{L}$ of this mix into the wells containing the corresponding experimental community matching each species pool. After this, the experimental cultures were thoroughly mixed using a 96-well pipettor (Viaflo 96, Integra Biosciences; settings: pipette/mix program, 5 mixing cycles, mixing volume 150 $\mu\text{L}$, speed 6) before applying a 30-fold dilution by transferring 10 $\mu\text{L}$ of the cultures into a new plate with 300 $\mu\text{L}$ of fresh media.

The resident communities were cultured over 6 dilution cycles before introducing the invader species. When introducing the invader species on day 6, The volume ratio between the monoculture of invader species and the resident community was $10^{-3}$. After introducing the invaders on day 6, we performed another 6 daily dilution cycles with a $10^{-5}$ dispersal rate for all species including the invader species until the end of the experiment. At the end of every daily cycle, 150uL samples of each culture were used to measure the OD (600nm), a proxy for the total biomass in the cultures, using a Varioskan Flash (Thermo Fisher Scientific) plate reader. The remaining culture volume was stored at -80 °C for subsequent DNA extraction.

##### DNA extraction, 16S rRNA sequencing and data analysis.

To monitor the dynamics of the microbial communities, we measured community composition via 16S ribosomal RNA (rRNA) amplicon sequencing. DNA extraction was performed by the Environmental Sample Preparation and Sequencing Facility at Argonne National Laboratory. The obtained DNA was used for 16S (V4 region) amplicon sequencing. Library preparation and Illumina MiSeq sequencing were performed by the Environmental Sample Preparation and Sequencing Facility at Argonne National Laboratory. We used the R package DADA2 to obtain the amplicon sequence variants (ASVs) as described by Callahan *et al.*<sup>1</sup>. Taxonomic identities were assigned to the ASVs by using SILVA (version 132) as a reference database. For each sample, species richness was calculated as the number of ASVs with a relative abundance $\geq 0.08\%$, which corresponds to the 0.08% extinction threshold used in simulation. Taxonomic identities were assigned to ASVs using Randomized Axelerated Maximum Likelihood (RaxML) using default parameters. In our sequencing dataset, the average sequencing depth is 17075 reads. This means that we could not effectively resolve any species abundance on the order of 0.01% or below. Our main observables, invasion success, diversity and stability, were calculated (Methods) only from species abundances that exceed a threshold of 0.08% (the extinction threshold).

##### Numerical methods

We modeled the long-term dynamics and diversity of ecological communities using the well-known generalized Lotka-Volterra (gLV) model, modified to include dispersal from a species pool:

$$\frac{dN_i}{dt} = r_i N_i \left( 1 - \sum_{j=1}^s \alpha_{ij} N_j / K_i \right) + D \quad (1)$$

where $N_i$ is the abundance of species $i$ (normalized to its carrying capacity), $\alpha_{ij}$ is the interaction strength that captures how strongly species $j$ inhibits the growth of species $i$ (with self-regulation $\alpha_{ii} = 1$), and $D$ is the dispersal rate from an outside species pool to the focal community. For simplicity and without qualitatively changing our results, we considered the same growth rate $r_i = 1$ and the same carrying capacity $K_i = 1$ for all species in the main text. Our previous paper shows that sampling growth rates from a uniform distribution has little effect on the phase diagram of survival fraction and fluctuation fraction<sup>2</sup>. Our previous paper also shows that sampling carrying capacities from a normal distribution increases the partial coexistence phase while shrinking both the full coexistence phase and fluctuation phase but does not affect the order of the phases<sup>2</sup>.

In our previous work, we tested the theoretical predictions when considering the existence of positive (facilitative) interspecies interactions and varying the symmetry of the interaction matrix<sup>2</sup>. We also considered different dispersal rates, and the effects of incorporating daily dilutions in these *in silico* communities<sup>2</sup>. These additional results show that our qualitative phase diagrams and conclusions are robust to different choices of ecological network structure and parameters. Although the patterns of ecological diversity and dynamics do not change as the dispersal rate varies from $D=10^{-7}$ to $D=10^{-6}$, we found that communities with zero dispersal rate exhibit lower fluctuation fraction and survival fraction in the persistent fluctuation phase<sup>2</sup>. Our results showed that non-zero dispersal rates can sustain persistent fluctuations. After the resident species typically reach steady states at $t=10^3$, we started introducing the invader species by continuously adding dispersal of the invader to the resident community and simulated the dynamics until $t=2\times 10^3$ to determine the invasion outcome.

All simulations used the Runge-Kutta method on Matlab to numerically solve the LV equations (with an integration step of 0.05). A definition of $20\times 20$ pixels was used for each phase diagram (Fig. 2f and 4c), linearly segmenting the parameter space in the ranges $\langle \alpha_{ij} \rangle \in [0.02, 1.1]$ and $S \in [2, 60]$. In each phase diagram, each pixel shows the average result for $10^3$ simulations. The total simulation time is $2\times 10^3$. We sampled the interaction strength from a uniform distribution $U[0.5\langle \alpha_{ij} \rangle, 1.5\langle \alpha_{ij} \rangle]$ in Fig. 2b and 4d, where $\langle \alpha_{ij} \rangle$ is the mean interaction strength between species (which also determines here the variance of interactions).

##### Reaching steady state, extinction threshold, survival fraction and stability in simulations

We define the steady state of simulated communities as the community state in which community properties (e.g., survival fraction, fluctuation fraction, and invasion probability) significantly changes as time goes on. To consistently analyze the steady state results for all the simulated communities, in our previous work, we analyzed the dependence of the phase diagrams on the simulated time<sup>2</sup>. Our results showed that community-level properties did not significantly changes after $t=10^3$. Accordingly, the phase diagrams in the paper show the state of communities at $t=2\times 10^3$, unless otherwise stated.

The presence of dispersal from the species pool in Eq. (1) guarantees that all species exhibit strictly positive abundances in Fig 2a. Nevertheless, we consider that a species is extinct if its abundance lays below an $8\times 10^{-4}$ threshold, as consistent with the extinction in our experiment. Around this threshold, the dispersal rate becomes the main factor preventing abundance decay<sup>2</sup>. The species abundance distribution in the partial coexistence phase is bimodal<sup>2</sup>; the extinction threshold $8\times 10^{-4}$ clearly separates the high-abundance surviving species from low-abundance species that will go extinct if dispersal ceases<sup>2</sup>.

To compute the survival fraction, we computed the fraction of species whose abundance exceeded the extinction threshold at any time during the last 100 units of time in the simulation. The survival or extinction of invader species was determined by the abundance in the last 100 units of time in the simulation. Our choice of including a time window when measuring diversity is motivated by the fact that, for the case of unstable communities, species abundances fluctuate above and below the extinction threshold over time. Since we measured diversity and species compositions every 24 hours in the experiment, we consider an analogous window of 100 time units in simulations.

To differentiate between stable and fluctuating communities, we computed the average coefficient of variation of $N_i$ between $t=10^3-100$ and $t=10^3$. We define communities with this average coefficient of variation of species abundance higher (lower) than $10^{-3}$ as fluctuating (stable) communities.

##### Definition of stable and fluctuating experimental communities

To differentiate between stable and fluctuating resident communities in experiments, we computed the standard deviation of biomass between day 4, day 5 and day 6. Communities for which the standard deviation of biomass over time is below (above) a 0.05 threshold are considered stable (fluctuating) communities (Supplementary Fig. 12). We also calculated the average coefficient of variation (CV) for species abundances from day 4 to day 6. This corresponds to the average value of the standard deviation for the absolute abundance of each species $N_i$ (over day 4, day 5, and day 6) scaled by average species abundance. The average coefficient of variation of absolute species abundance (product of biomass and relative species abundance by 16s sequencing) displays a strong positive correlation with the standard deviation of biomass over time across communities (Supplementary Fig. 12). The average coefficient of variation of relative species abundance (by 16s sequencing) also displays a strong positive correlation with the standard deviation of biomass over time across communities (Supplementary Fig. 12). Different metrics consistently classify the communities into two clusters: fluctuating ones on top right region and stable ones on bottom left region (Supplementary Fig. 12). Varying the choice of time window (day 4 to day 6) to a new time window (day 5 to day 6) yield the same classification of fluctuating and stable communities. Our previous work showed that the classification of stability converges quickly to either small or large values, respectively indicating stability or long-lasting fluctuations in experimental communities<sup>2</sup>. We found the $K$-means clustering algorithm yields the same classification results of community stability. The consistence between results of $K$-means clustering and setting stability threshold of biomass standard deviation (0.05) demonstrates the classification of fluctuating and stable communities is robust to different algorithm.

● Bacteria-Firmicutes-Bacilli-Lactobacillales-Streptococcaceae-Lactococcus
● Bacteria-Proteobacteria-Gammaproteobacteria-Enterobacterales-Enterobacteriaceae-Raoultella
● Bacteria-Proteobacteria-Gammaproteobacteria-Enterobacterales-Enterobacteriaceae-Klebsiella
● Bacteria-Bacteroidota-Bacteroidia-Flavobacteriales-Weeksellaceae-Chryseobacterium
● Bacteria-Proteobacteria-Gammaproteobacteria-Enterobacterales-Enterobacteriaceae-Pluralibacter
● Bacteria-Firmicutes-Bacilli-Lactobacillales-Leuconostocaceae-Leuconostoc
● Bacteria-Proteobacteria-Gammaproteobacteria-Aeromonadales-Aeromonadaceae-Aeromonas
● Bacteria-Proteobacteria-Gammaproteobacteria-Enterobacterales-NA-NA
● Bacteria-Bacteroidota-Bacteroidia-Flavobacteriales-Weeksellaceae-Empedobacter
● Bacteria-Proteobacteria-Gammaproteobacteria-Xanthomonadales-Xanthomonadaceae-Stenotrophomonas
● Bacteria-Bacteroidota-Bacteroidia-Flavobacteriales-Weeksellaceae-Empedobacter
● Bacteria-Bacteroidota-Bacteroidia-Sphingobacteriales-Sphingobacteriaceae-Sphingobacterium
● Bacteria-Proteobacteria-Gammaproteobacteria-Enterobacterales-Erwiniaaceae-Pantoea
● Bacteria-Firmicutes-Bacilli-Lactobacillales-Leuconostocaceae-Leuconostoc
● Bacteria-Proteobacteria-Alphaproteobacteria-Rhizobiales-Rhizobiaceae-Ochrobactrum
● Bacteria-Proteobacteria-Gammaproteobacteria-Pseudomonadales-Pseudomonadaceae-Pseudomonas
● Bacteria-Firmicutes-Bacilli-Exiguobacterales-Exiguobacteraceae-Exiguobacterium
● Bacteria-Proteobacteria-Gammaproteobacteria-Enterobacterales-Enterobacteriaceae-Escherichia/Shigella
● Bacteria-Firmicutes-Bacilli-Bacillales-Planococcaceae-Lysinibacillus
● Bacteria-Proteobacteria-Gammaproteobacteria-Pseudomonadales-Moraxellaceae-Acinetobacter
● Bacteria-Bacteroidota-Bacteroidia-Flavobacteriales-Weeksellaceae-Empedobacter
● Bacteria-Bacteroidota-Bacteroidia-Flavobacteriales-Weeksellaceae-Empedobacter
● Bacteria-Firmicutes-Bacilli-Staphylococcales-Staphylococcaceae-Staphylococcus
● Bacteria-Proteobacteria-Gammaproteobacteria-Pseudomonadales-Pseudomonadaceae-Pseudomonas
● Bacteria-Bacteroidota-Bacteroidia-Sphingobacteriales-Sphingobacteriaceae-Pedobacter
● Bacteria-Bacteroidota-Bacteroidia-Cytophagales-Spirosomaceae-Flectobacillus
● Bacteria-Proteobacteria-Gammaproteobacteria-Burkholderiales-Oxalobacteraceae-Herbaspirillum
● Bacteria-Firmicutes-Bacilli-Bacillales-Bacillaceae-Bacillus
● Bacteria-Proteobacteria-Gammaproteobacteria-Pseudomonadales-Pseudomonadaceae-Pseudomonas
● Bacteria-Proteobacteria-Gammaproteobacteria-Xanthomonadales-Xanthomonadaceae-Stenotrophomonas
● Bacteria-Proteobacteria-Gammaproteobacteria-Burkholderiales-Oxalobacteraceae-Undibacterium
● Bacteria-Proteobacteria-Gammaproteobacteria-Enterobacterales-Enterobacteriaceae-NA
● Bacteria-Proteobacteria-Gammaproteobacteria-Enterobacterales-Enterobacteriaceae-Raoultella
● Bacteria-Bacteroidota-Bacteroidia-Cytophagales-Spirosomaceae-Flectobacillus
● Bacteria-Proteobacteria-Gammaproteobacteria-Aeromonadales-Aeromonadaceae-Aeromonas
● Bacteria-Proteobacteria-Gammaproteobacteria-Burkholderiales-Comamonadaceae-Acidovorax
● Bacteria-Bacteroidota-Bacteroidia-Flavobacteriales-Weeksellaceae-Chryseobacterium
● Bacteria-Proteobacteria-Gammaproteobacteria-Enterobacterales-Enterobacteriaceae-Citrobacter
● Bacteria-Firmicutes-Bacilli-Lactobacillales-Streptococcaceae-Lactococcus
● Bacteria-Firmicutes-Bacilli-Bacillales-Planococcaceae-NA
● Bacteria-Proteobacteria-Gammaproteobacteria-Enterobacterales-Erwiniaaceae-Pantoea
● Bacteria-Proteobacteria-Gammaproteobacteria-Pseudomonadales-Pseudomonadaceae-Pseudomonas
● Bacteria-Proteobacteria-Gammaproteobacteria-Enterobacterales-Enterobacteriaceae-Klebsiella
● Bacteria-Proteobacteria-Gammaproteobacteria-Enterobacterales-Enterobacteriaceae-Enterobacter
● Bacteria-Proteobacteria-Gammaproteobacteria-Pseudomonadales-Pseudomonadaceae-Pseudomonas
● Bacteria-Firmicutes-Bacilli-Staphylococcales-Staphylococcaceae-Staphylococcus
● Bacteria-Firmicutes-Bacilli-Bacillales-Bacillaceae-Bacillus
● Bacteria-Firmicutes-Bacilli-Exiguobacterales-Exiguobacteraceae-Exiguobacterium
● Bacteria-Bacteroidota-Bacteroidia-Flavobacteriales-Flavobacteriaceae-Flavobacterium
● Bacteria-Proteobacteria-Gammaproteobacteria-Pseudomonadales-Pseudomonadaceae-Pseudomonas
● Bacteria-Proteobacteria-Gammaproteobacteria-Aeromonadales-Aeromonadaceae-Aeromonas
● Bacteria-Proteobacteria-Gammaproteobacteria-Aeromonadales-Aeromonadaceae-Tolomonas
● Bacteria-Proteobacteria-Gammaproteobacteria-Enterobacterales-NA-NA
● Bacteria-Proteobacteria-Gammaproteobacteria-Enterobacterales-Enterobacteriaceae-Pluralibacter
● Bacteria-Actinobacteriota-Actinobacteria-Streptomycetales-Streptomycetaceae-Streptomyces
● Bacteria-Actinobacteriota-Actinobacteria-Micrococcales-Microbacteriaceae-Curtobacterium
● Bacteria-Proteobacteria-Gammaproteobacteria-Pseudomonadales-Pseudomonadaceae-Pseudomonas
● Bacteria-Firmicutes-Bacilli-Lactobacillales-Leuconostocaceae-Leuconostoc
● Bacteria-Bacteroidota-Bacteroidia-Bacteroidales-Williamwhitmaniaceae-Acetobacteroides
● Bacteria-Proteobacteria-Gammaproteobacteria-Xanthomonadales-Xanthomonadaceae-Stenotrophomonas
● Bacteria-Proteobacteria-Gammaproteobacteria-Aeromonadales-Aeromonadaceae-Tolomonas
● Bacteria-Bacteroidota-Bacteroidia-Flavobacteriales-Weeksellaceae-Empedobacter
● Bacteria-Firmicutes-Bacilli-Exiguobacterales-Exiguobacteraceae-Exiguobacterium
● Bacteria-Bacteroidota-Bacteroidia-Flavobacteriales-Weeksellaceae-Empedobacter
● Bacteria-Proteobacteria-Gammaproteobacteria-Enterobacterales-Erwiniaaceae-Pantoea
● Bacteria-Firmicutes-Bacilli-Bacillales-Bacillaceae-Bacillus
● Bacteria-Proteobacteria-Gammaproteobacteria-Burkholderiales-Oxalobacteraceae-Undibacterium
● Bacteria-Proteobacteria-Gammaproteobacteria-Pseudomonadales-Moraxellaceae-Acinetobacter
● Bacteria-Proteobacteria-Gammaproteobacteria-Enterobacterales-Enterobacteriaceae-Citrobacter
● Bacteria-Proteobacteria-Gammaproteobacteria-Enterobacterales-Enterobacteriaceae-Escherichia/Shigella
● Bacteria-Firmicutes-Bacilli-Lactobacillales-Streptococcaceae-Lactococcus
● Bacteria-Bacteroidota-Bacteroidia-Flavobacteriales-Flavobacteriaceae-Flavobacterium
● Bacteria-Proteobacteria-Alphaproteobacteria-Rhizobiales-Rhizobiaceae-Ochrobactrum
● Bacteria-Firmicutes-Clostridia-Lachnospirales-Lachnospiraceae-Lachnospiraceae\_NK4A136\_group
● Bacteria-Cyanobacteria-Cyanobacteriia-Chloroplast-NA-NA
● Bacteria-Firmicutes-Bacilli-Bacillales-Planococcaceae-Lysinibacillus
● Bacteria-Proteobacteria-Gammaproteobacteria-Enterobacterales-Enterobacteriaceae-NA
● Bacteria-Proteobacteria-Gammaproteobacteria-Burkholderiales-Comamonadaceae-Acidovorax
● Bacteria-Bacteroidota-Bacteroidia-Flavobacteriales-Weeksellaceae-Chryseobacterium
● Bacteria-Firmicutes-Clostridia-Lachnospirales-Lachnospiraceae-Agathobacter

**Supplementary Fig. 1. Taxonomic identity of the bacterial isolates.** The identities have been inferred from the ASV (Methods) of 16S sequencing, which allow the classification of the 80 isolates down to the genus level. Colors are consistent with those in the main text and other supplementary figures.

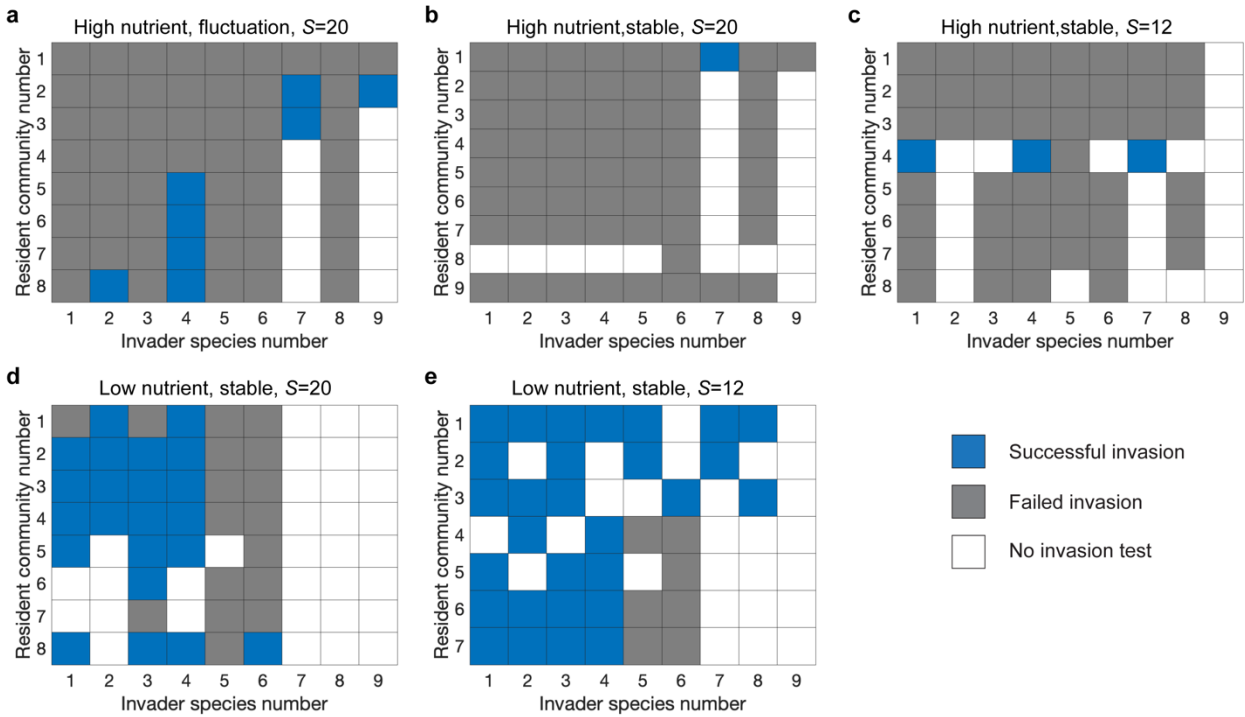

**Supplementary Fig. 2. Introducing different invaders into different resident communities and measuring the invasion outcome through 16s sequencing.** The invasion outcome matrices show that increasing nutrient and species pool size lead to a decrease in invasion probability.

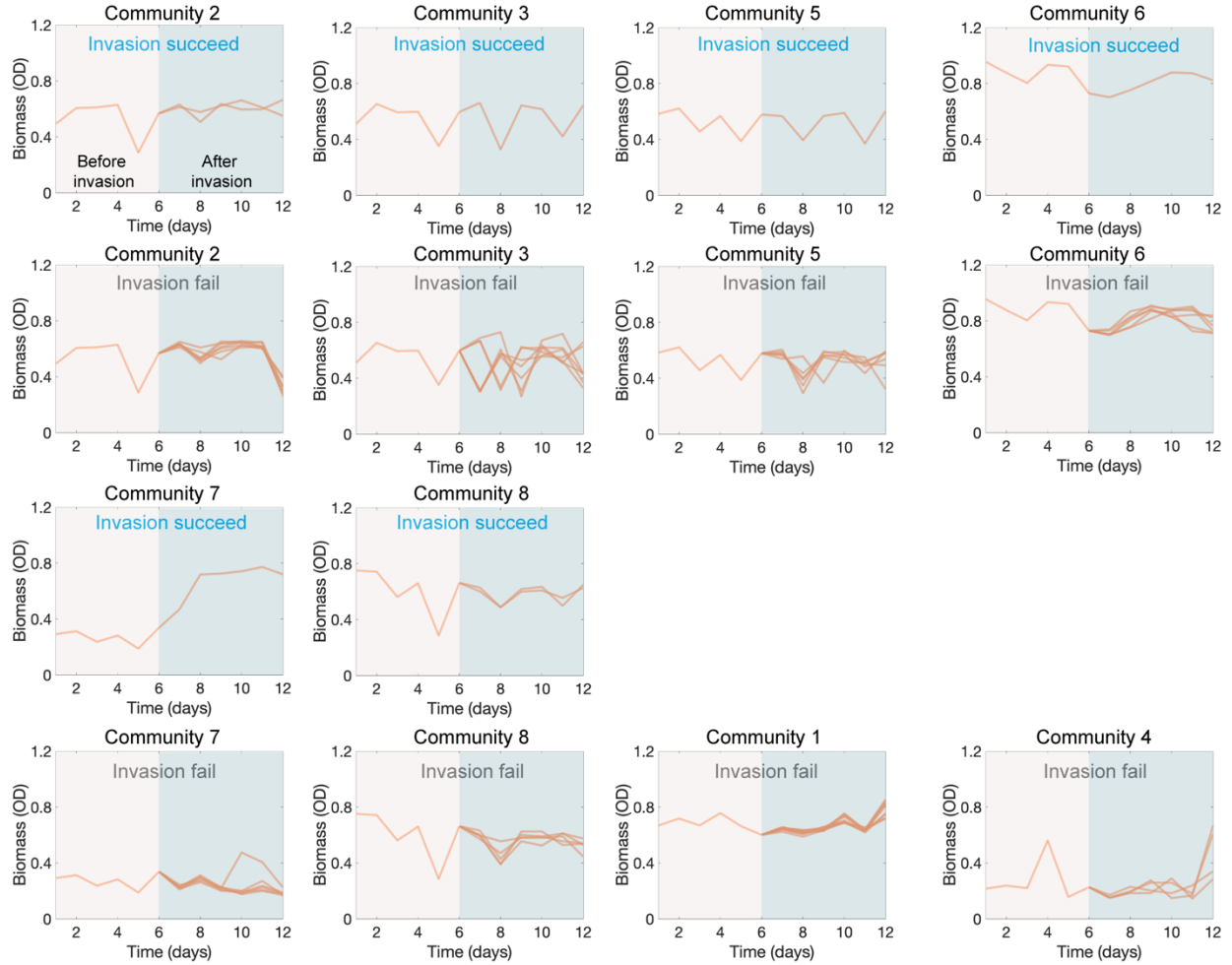

**Supplementary Fig. 3. Time series for the biomass of the fluctuating communities with species pool size $S=20$ under strong average interaction strength (high nutrients concentration).** Each panel shows the time series for the OD (600nm) of one fluctuating community with species pool size $S=20$ under high nutrient. The invaders were introduced on day 6, and the time series of successful invasions and failed invasions for the same communities were displayed in different panels.

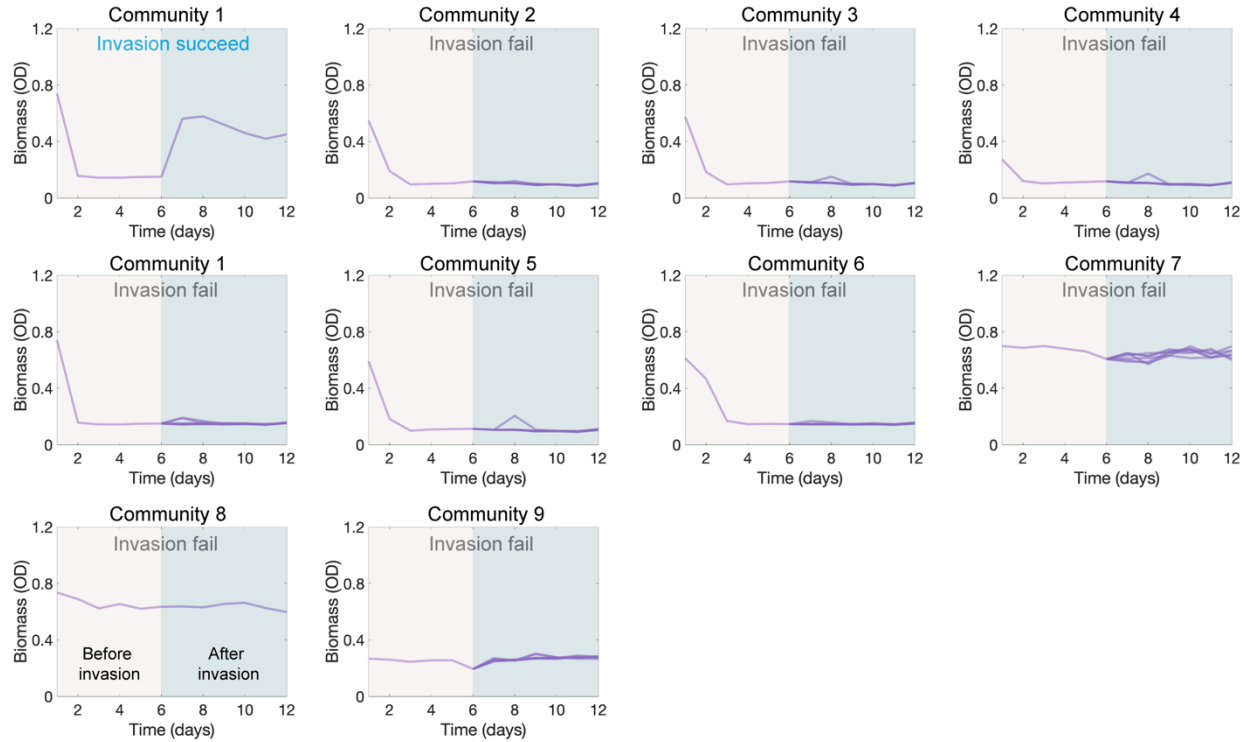

**Supplementary Fig. 4. Time series for the biomass of the stable communities with species pool size $S=20$ under strong average interaction strength (high nutrients concentration).** Each panel shows the time series for the OD (600nm) of one stable community with species pool size $S=20$ under high nutrient. The invaders were introduced on day 6, and the time series of successful invasions and failed invasions for the same communities were displayed in different panels.

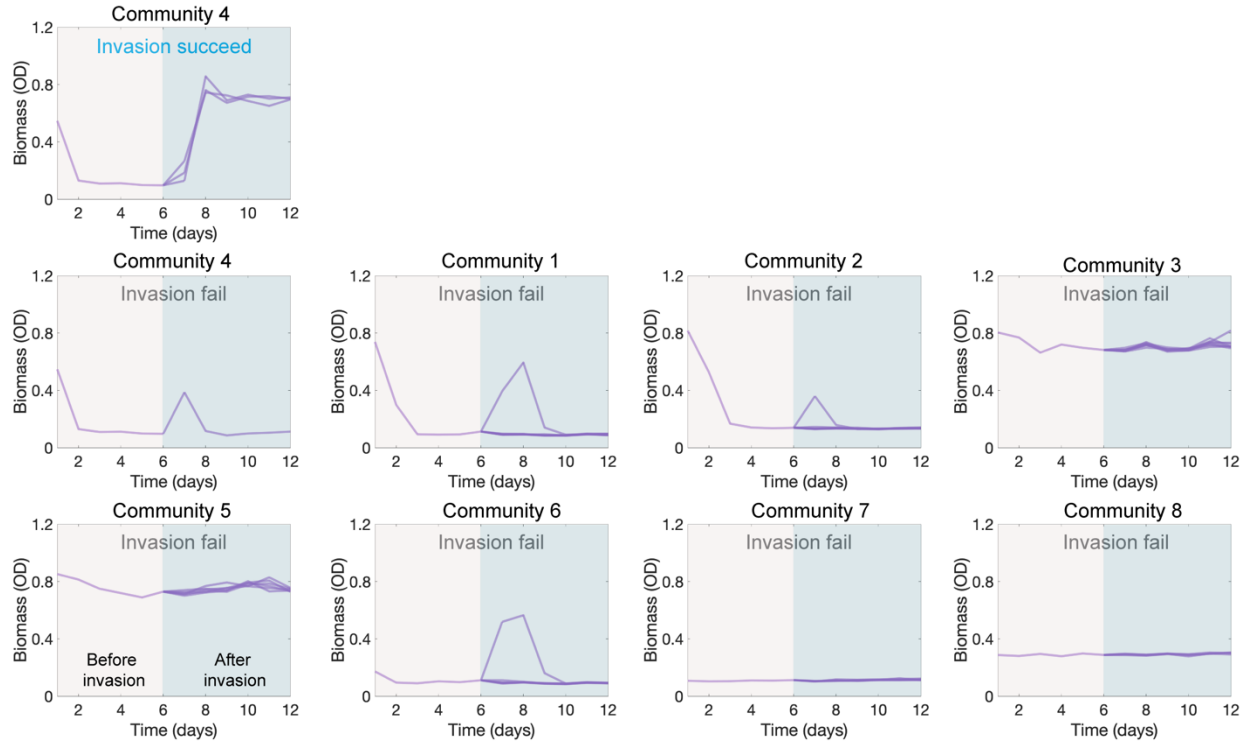

**Supplementary Fig. 5. Time series for the biomass of the stable communities with species pool size $S=12$ under strong average interaction strength (high nutrients concentration).** Each panel shows the time series for the OD (600nm) of one stable community with species pool size $S=12$ under high nutrient. The invaders were introduced on day 6, and the time series of successful invasions and failed invasions for the same communities were displayed in different panels.

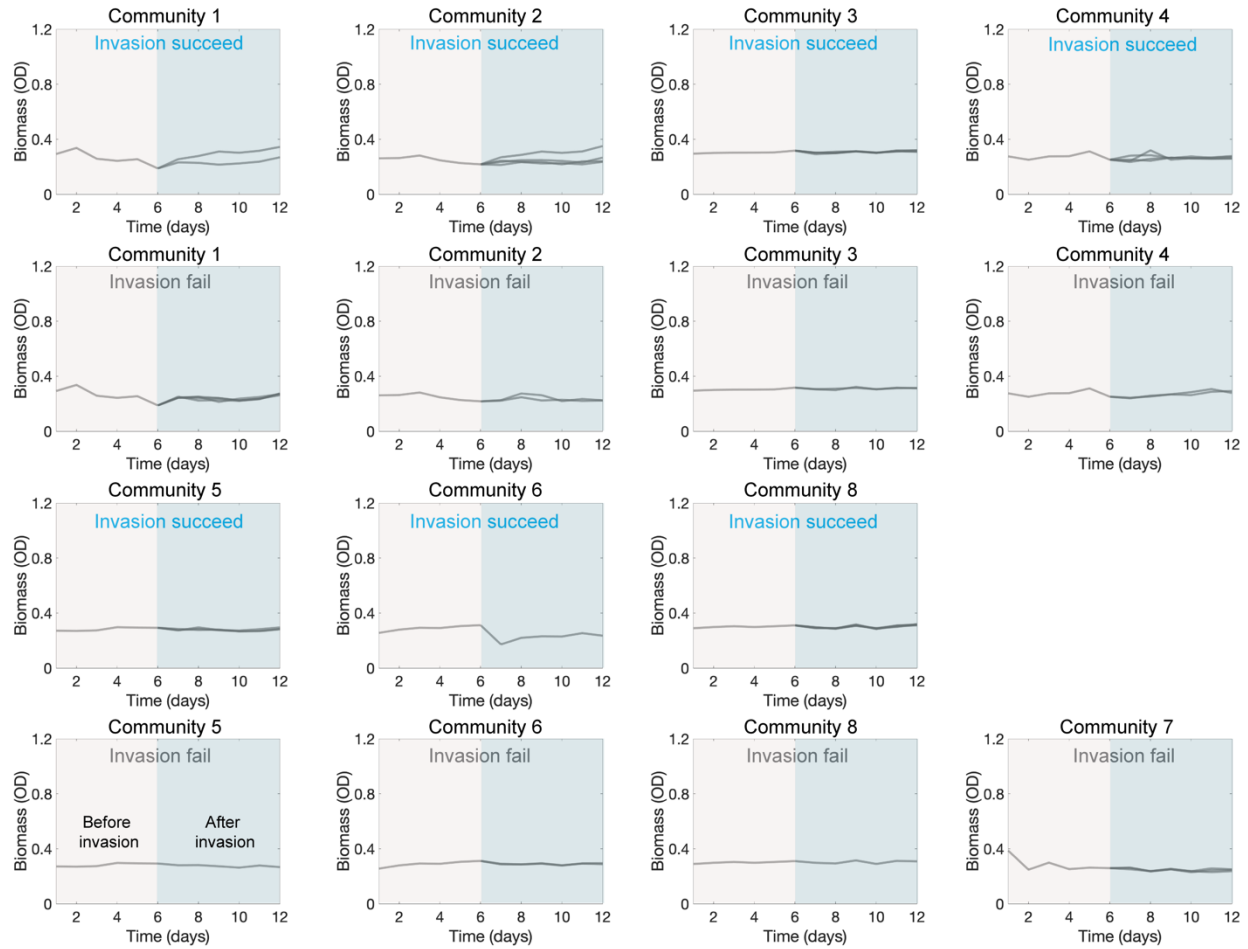

**Supplementary Fig. 6. Time series for the biomass of the stable communities with species pool size $S=20$ under weak average interaction strength (low nutrients concentration).** Each panel shows the time series for the OD (600nm) of one stable community with species pool size $S=20$ under low nutrient. The invaders were introduced on day 6, and the time series of successful invasions and failed invasions for the same communities were displayed in different panels.

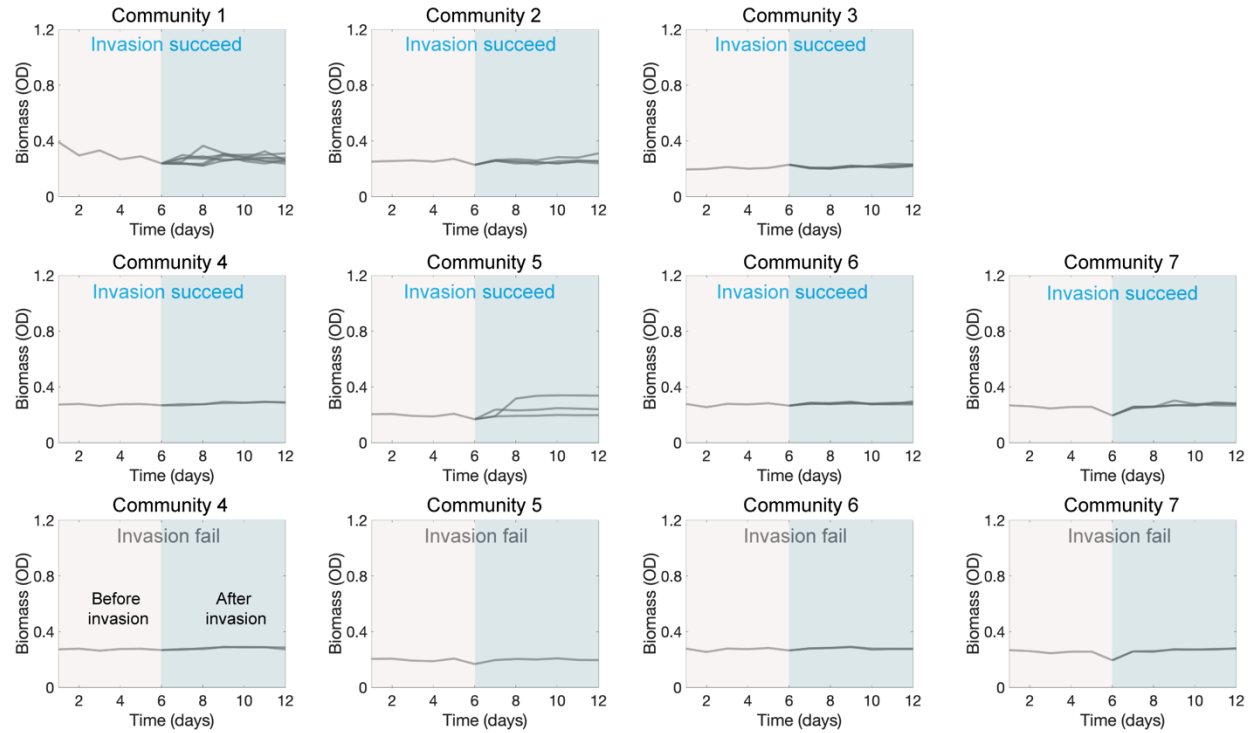

**Supplementary Fig. 7. Time series for the biomass of the stable communities with species pool size $S=12$ under weak average interaction strength (low nutrients concentration).** Each panel shows the time series for the OD (600nm) of one stable community with species pool size $S=12$ under low nutrient. The invaders were introduced on day 6, and the time series of successful invasions and failed invasions for the same communities were displayed in different panels.

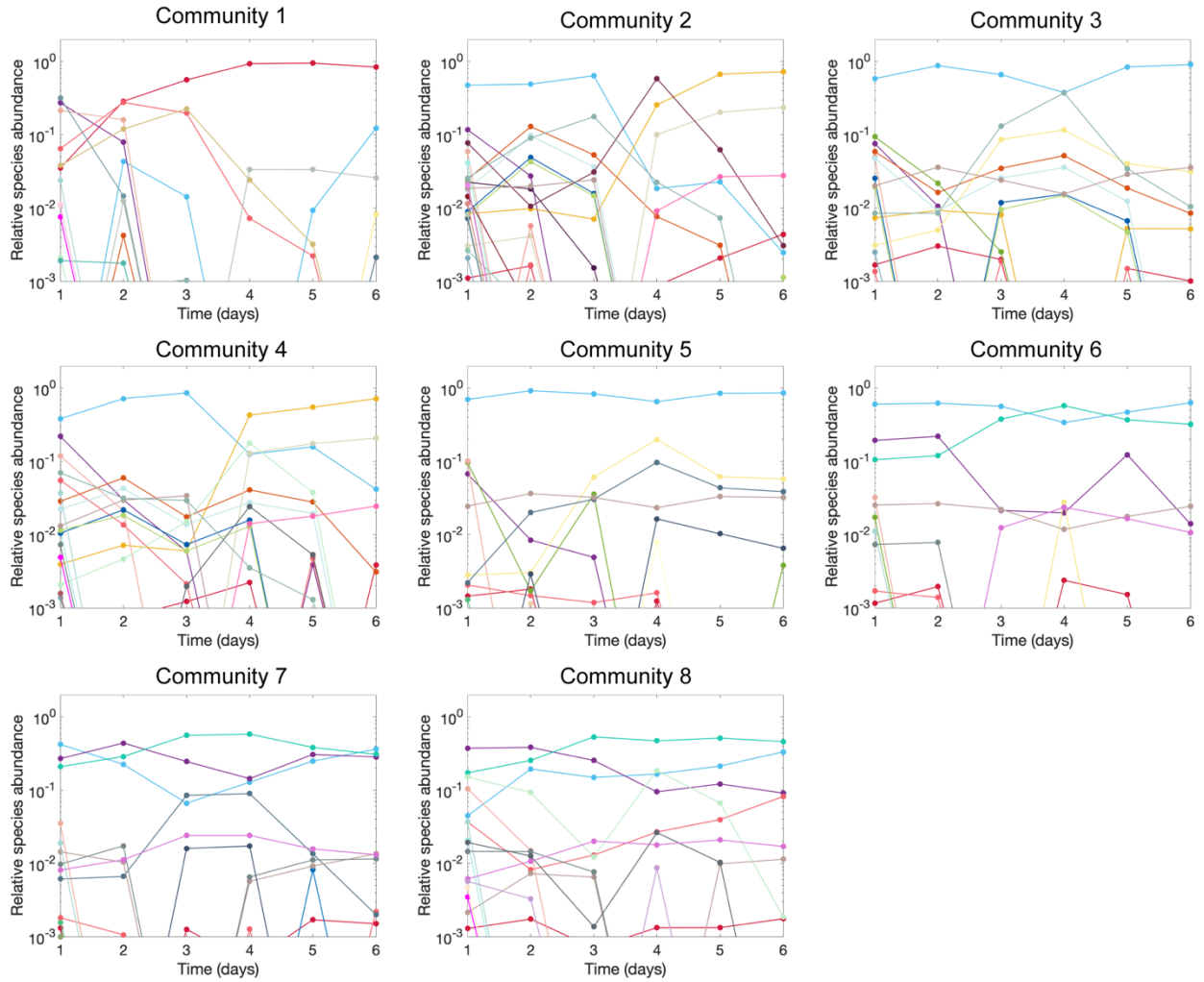

**Supplementary Fig. 8. Time series for the relative species abundances of the fluctuating communities with species pool size $S=20$ under strong average interaction strength (high nutrients concentration).** Each panel shows the time series for the relative species abundances of one fluctuating community before introducing invaders, where species pool size $S=20$ under high nutrient.

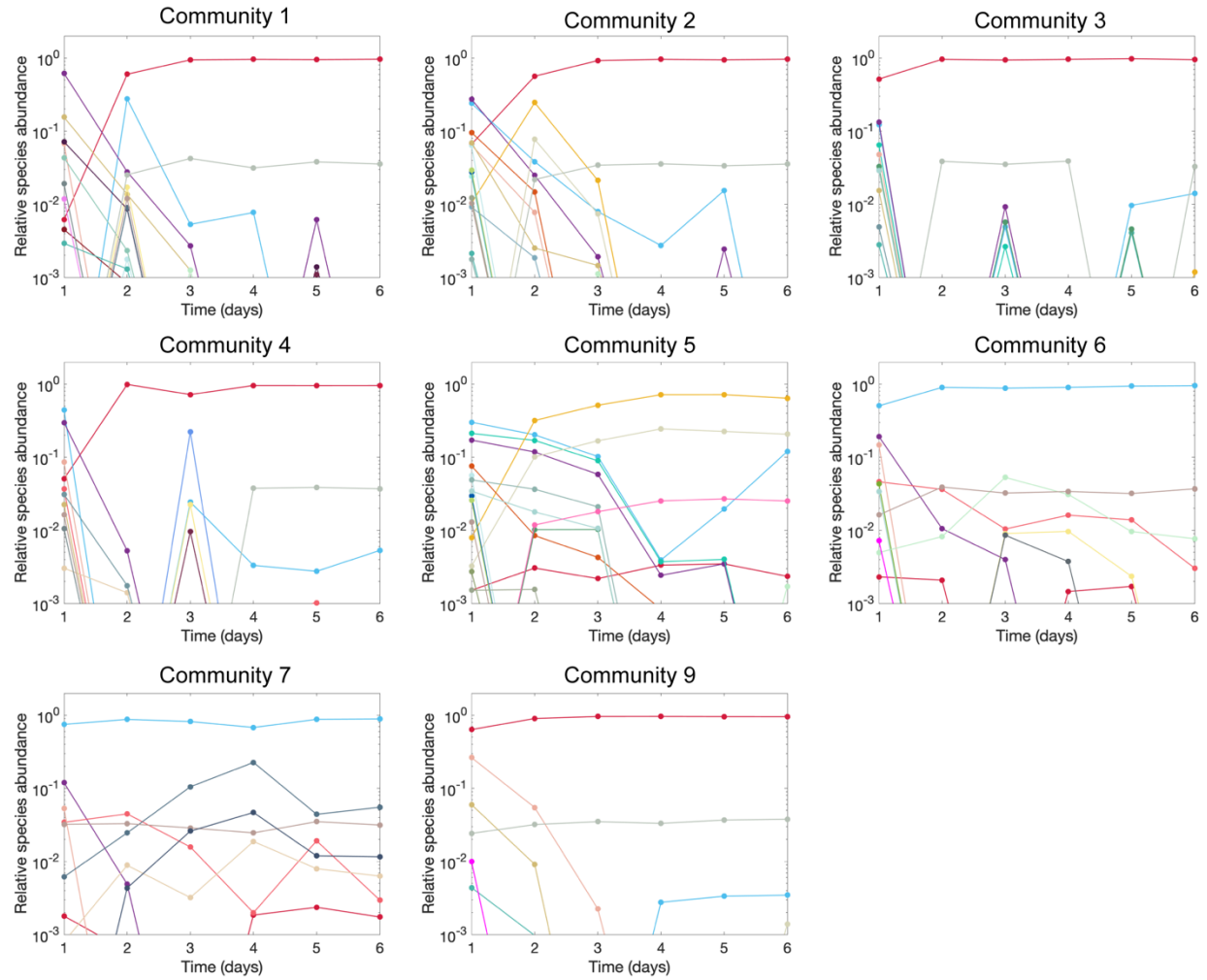

**Supplementary Fig. 9. Time series for the relative species abundances of the stable communities with species pool size $S=20$ under strong average interaction strength (high nutrients concentration). Each panel shows the time series for the relative species abundances of one stable community before introducing invaders, where species pool size $S=20$ under high nutrient.**

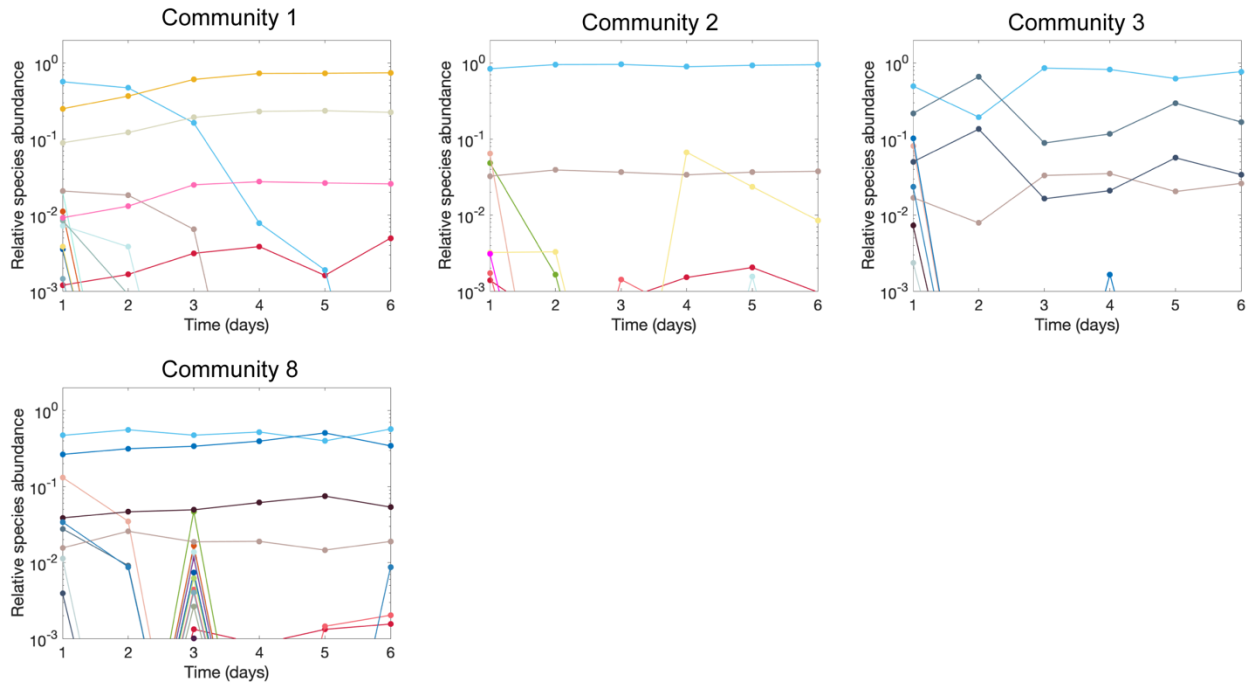

**Supplementary Fig. 10. Time series for the relative species abundances of the stable communities with species pool size $S=12$ under strong average interaction strength (high nutrients concentration).** Each panel shows the time series for the relative species abundances of one stable community before introducing invaders, where species pool size $S=12$ under high nutrient.

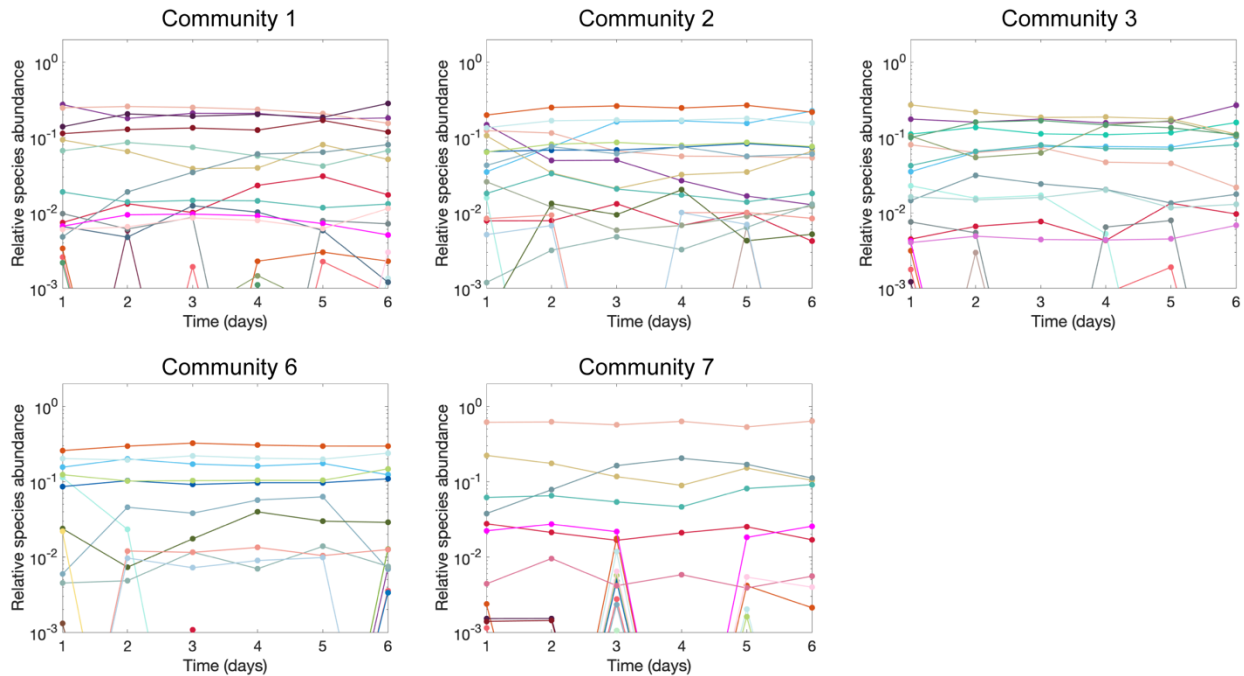

**Supplementary Fig. 11. Time series for the relative species abundances of the stable communities with species pool size $S=20$ under weak average interaction strength (low nutrients concentration).** Each panel shows the time series for the relative species abundances of one community before introducing invaders, where species pool size $S=20$ under low nutrient.

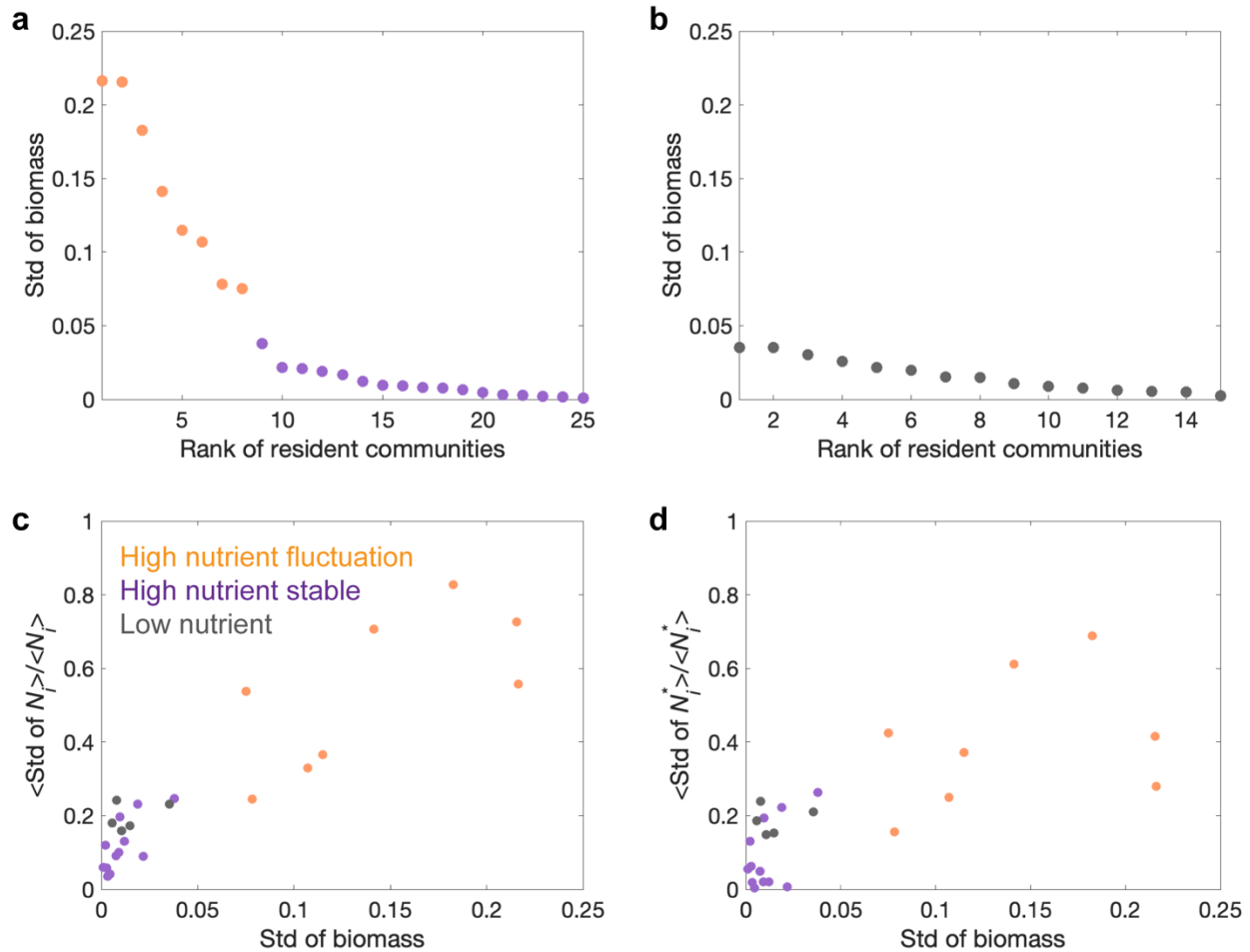

**Supplementary Fig. 12. Classification of fluctuating and stable resident communities in experiment.** **a**, The standard deviation of community biomass over day 5, day 6 and day 7 show that the stability threshold of 0.05 can separate the communities into stable ones (purple points) with small biomass deviation and fluctuating ones (orange points) with relatively large biomass deviation under high nutrient. **b**, The standard deviation of community biomass under low nutrient are small (all below the stability threshold of 0.05), which were naturally classified into stable communities. **c**, The average coefficient of (temporal) variation for absolute species abundances ($N_i$, computed as the product of total biomass per species relative abundance) exhibit a strong positive correlation with standard deviation of biomass in the experimental communities. The points span into two clusters where fluctuating communities locate on top right region and stable communities locate on bottom left region. **d**, The average coefficient of (temporal) variation for relative species abundances ($N_i^*$, relative species abundance through 16s sequencing) also exhibits a strong positive correlation with standard deviation of biomass in the experimental communities. The results suggest that fluctuation in community biomass cooccurs with fluctuation in relative species abundances.

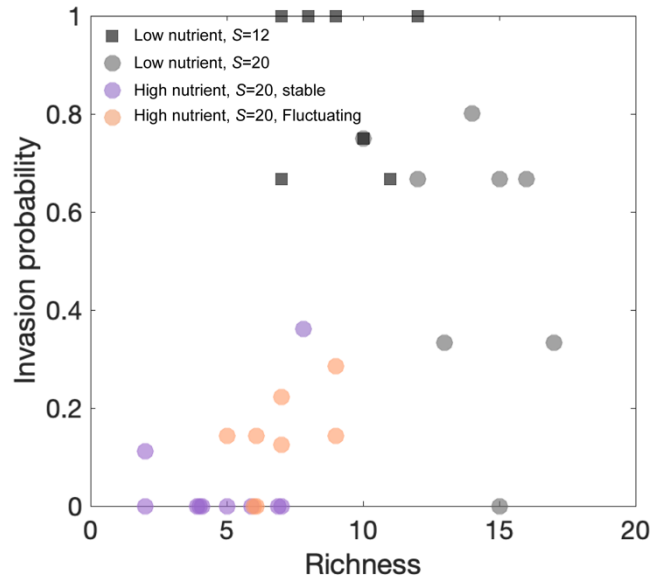

**Supplementary Fig. 13. Different invasibility-richness relationships in experiment depending upon how the richness is changed.** Invasibility positively correlates with richness when varying interaction strength (positive correlation between $S=20$ communities under low and high nutrient). Invasibility positively correlates with richness when randomly sample $S=20$ communities under high nutrient, due to fluctuating communities display larger richness and larger invasion probability. Invasibility negatively correlates with richness when increasing species pool size from $S=12$ to $S=20$ under low nutrient.

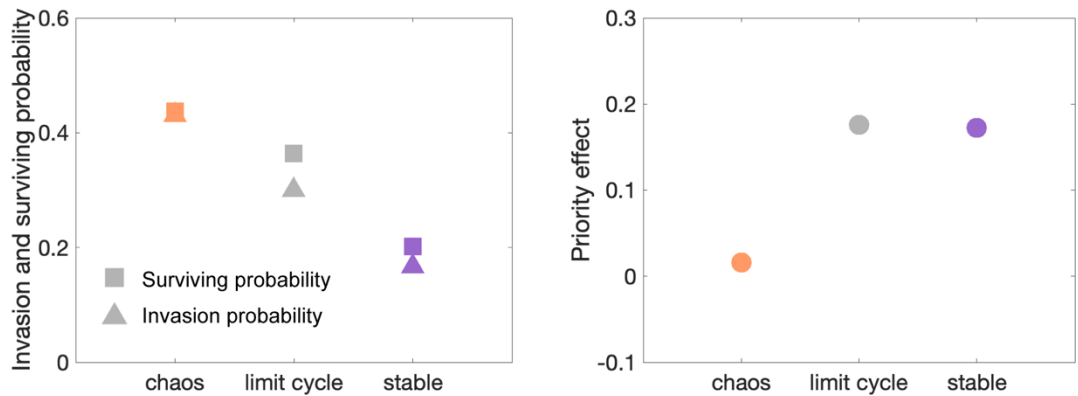

**Supplementary Fig. 14. Priority effect originates from alternative stable states and limit cycle oscillations rather than chaotic fluctuations in simulations.** Lotka-Volterra model simulations show that both surviving probability and invasion probability increase as community dynamics transition from alternative stable states to limit cycle oscillations and to chaos. Communities with chaotic fluctuations in species abundance do not display significant priority effect which can be explained by its ergodicity<sup>3,4</sup>, whereas communities with limit cycle oscillations and alternative stable states both show significant priority effect. The simulation in this figure was performed under $S=40$ and $\langle \alpha_{ij} \rangle = 0.65$ over 1000 replicates, among which we observed 223 chaotic fluctuating communities, 340 limit cycle oscillations, and 437 alternative stable states. The fluctuating communities were classified into chaos when its Lyapunov exponent is positive, while classified into limit cycle when its Lyapunov exponent is negative.

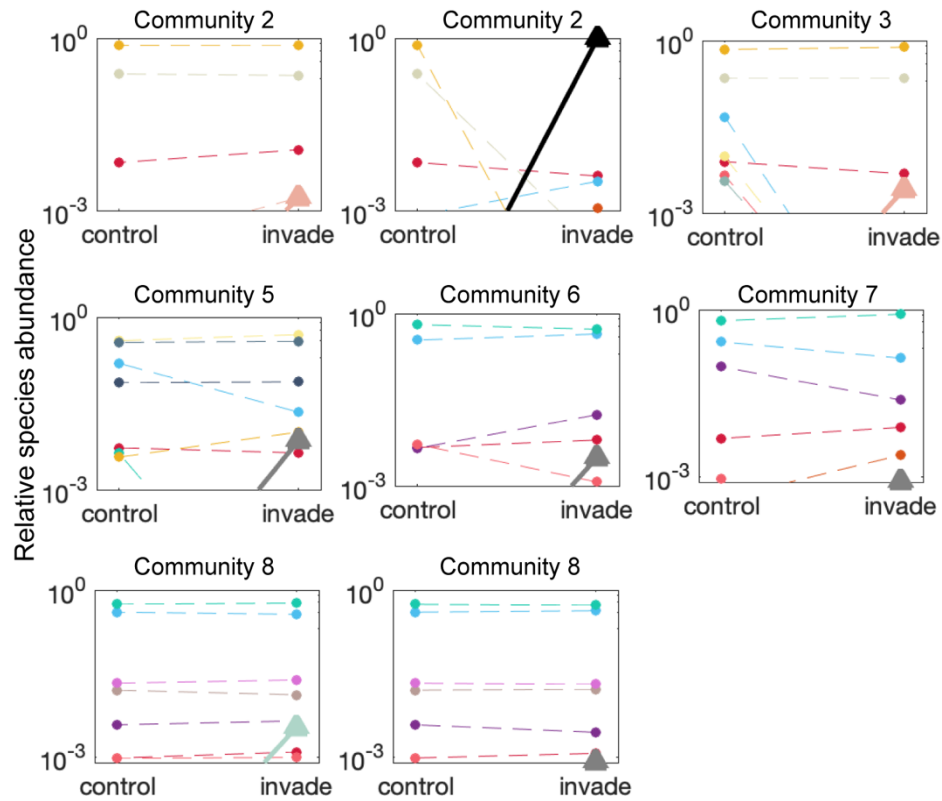

**Supplementary Fig. 15. Successful invasions lead to change in species composition in fluctuating communities with $S=20$ under high nutrient, which can be shown by comparing the relative species abundance between invaded communities and control communities without introducing invader.** The circles and triangles in the figure represent resident species and invader species, respectively. The successful invasions can cause the extinction of other resident species (circles drop below the extinction threshold under invasion) and the colonization of other resident species (circles go beyond the extinction threshold under invasion).

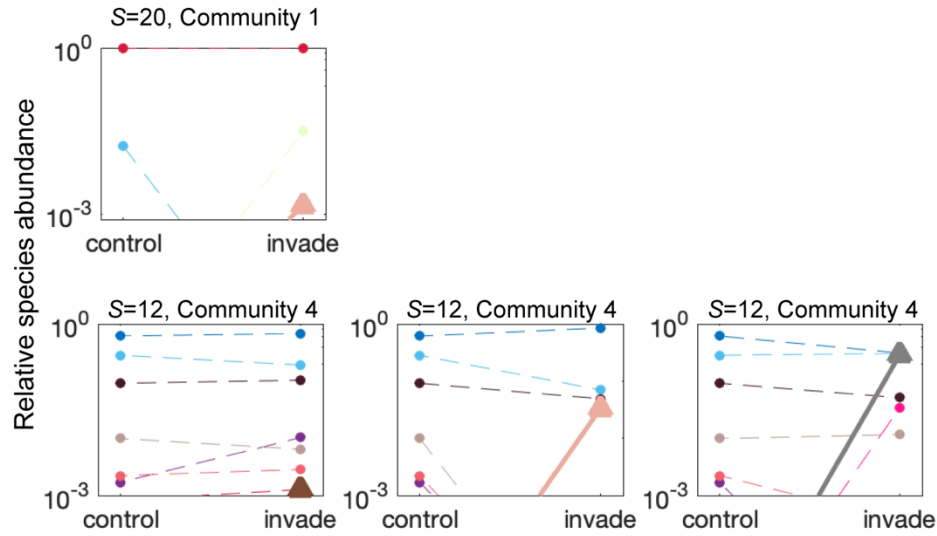

**Supplementary Fig. 16. Successful invasions lead to change in species composition in stable communities under high nutrient, which can be shown by comparing the relative species abundance between invaded communities and control communities without introducing invader.** The circles and triangles in the figure represent resident species and invader species, respectively. The successful invasions can cause the extinction of other resident species (circles drop below the extinction threshold under invasion) and the colonization of other resident species (circles go beyond the extinction threshold under invasion).

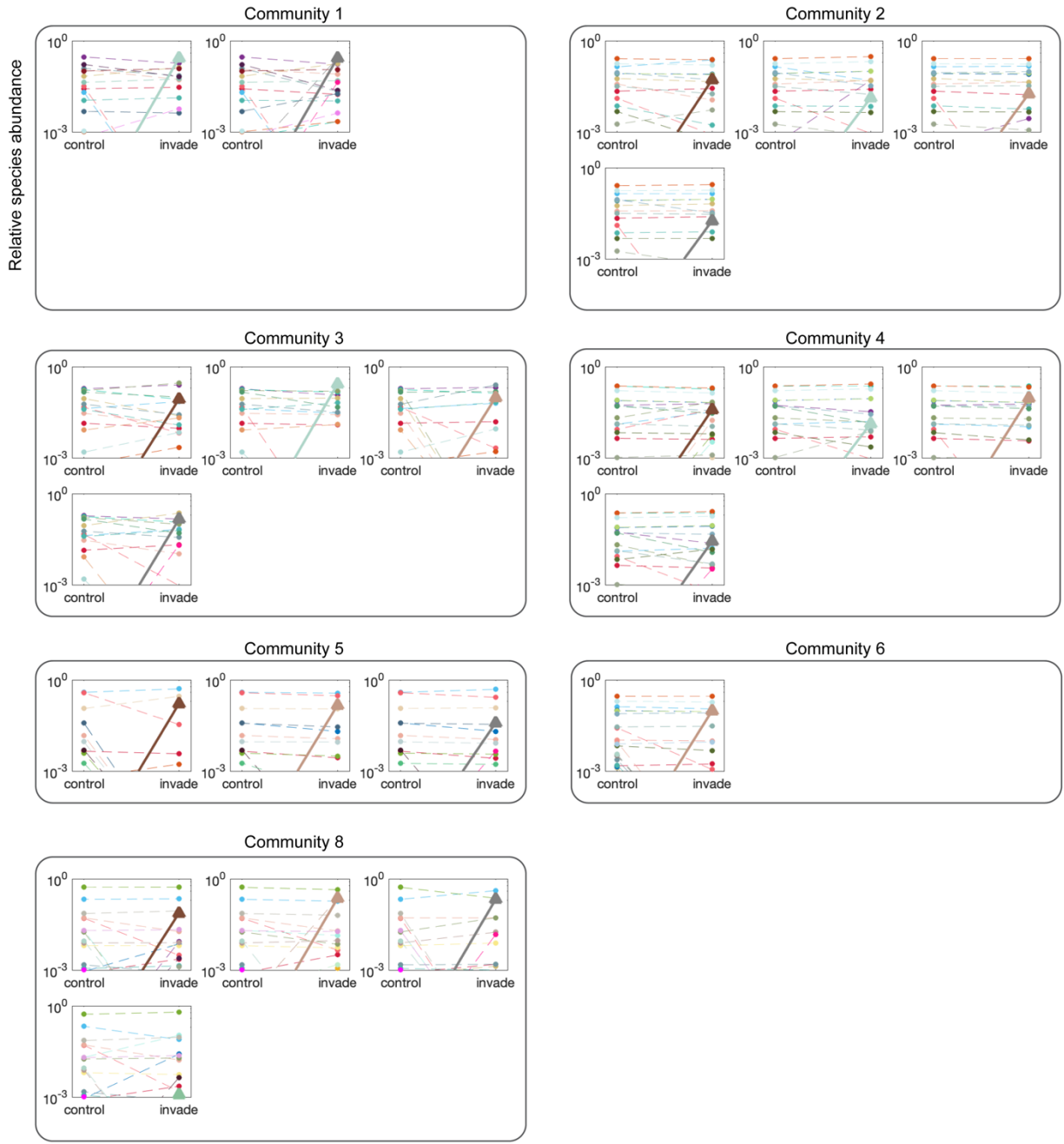

**Supplementary Fig. 17. Successful invasions lead to change in species composition in communities with $S=20$ under low nutrient, which can be shown by comparing the relative species abundance between invaded communities and control communities without introducing invader.** The circles and triangles in the figure represent resident species and invader species, respectively. The successful invasions can cause the extinction of other resident species (circles drop below the extinction threshold under invasion) and the colonization of other resident species (circles go beyond the extinction threshold under invasion).

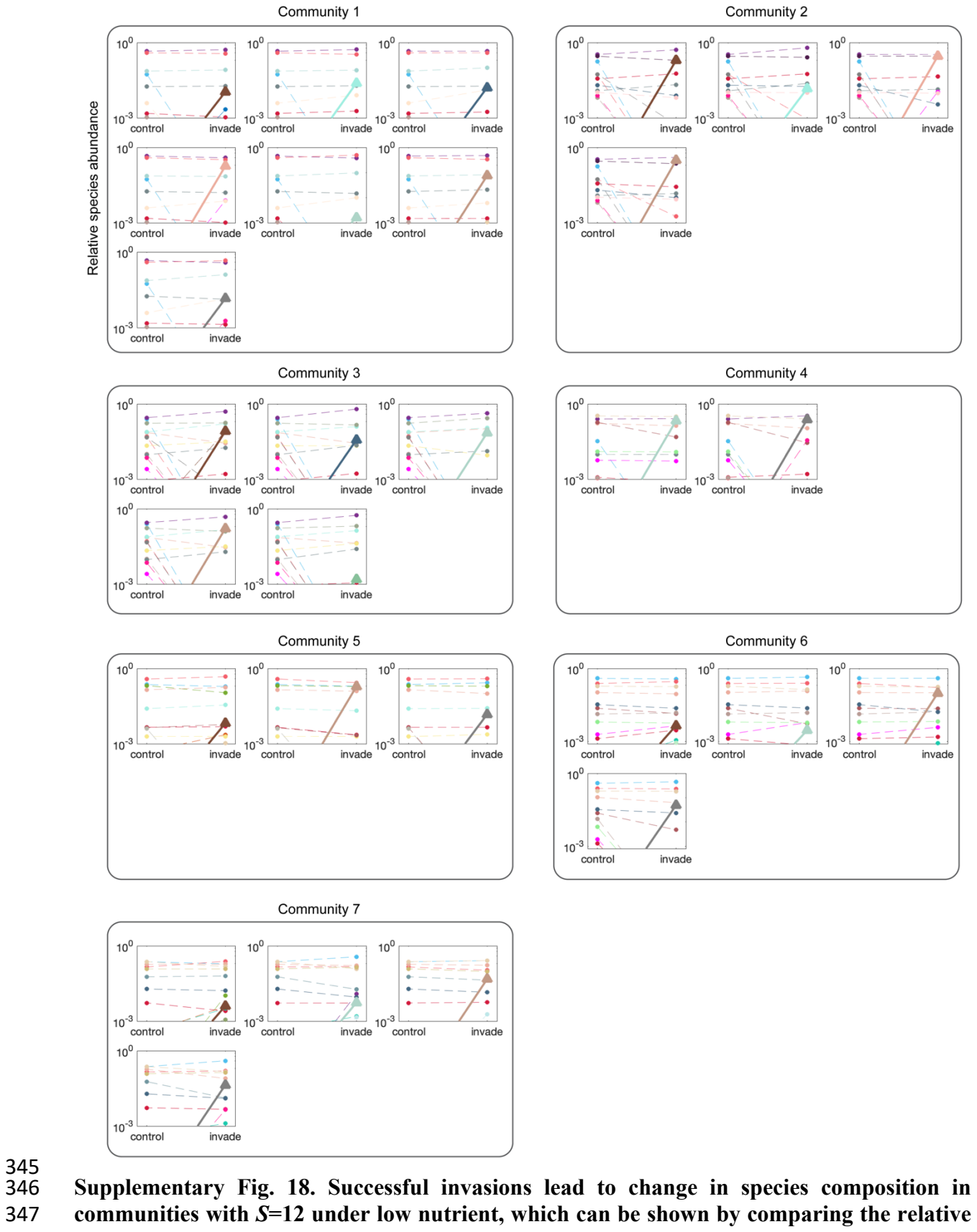

**species abundance between invaded communities and control communities without introducing invader.** The circles and triangles in the figure represent resident species and invader species, respectively. The successful invasions can cause the extinction of other resident species (circles drop below the extinction threshold under invasion) and the colonization of other resident species (circles go beyond the extinction threshold under invasion).

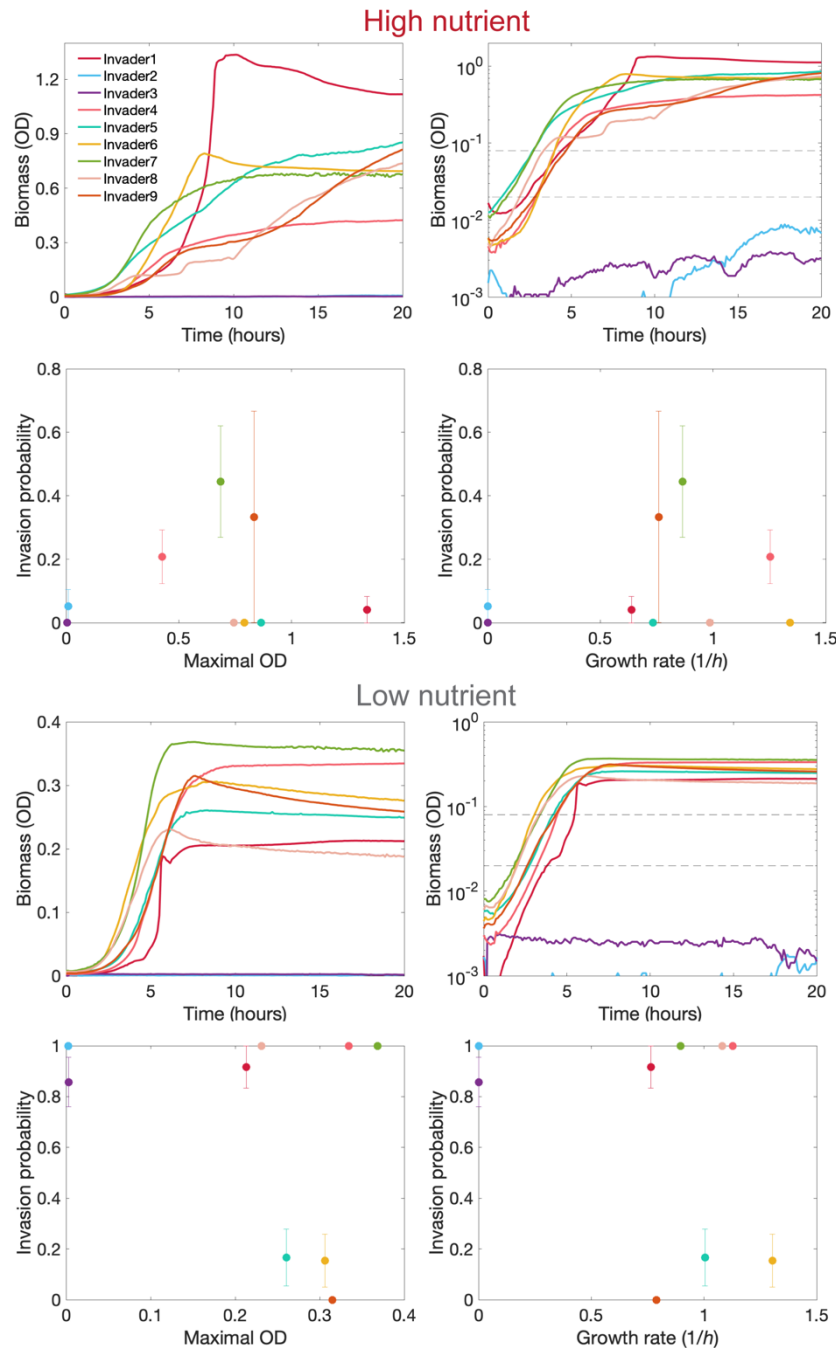

**Supplementary Fig. 19. There is no correlation between the invasiveness of invaders and their carrying capacities and growth rates.** The growth curves of invaders were measured after a dilution of $10^5$ folds. The carrying capacity of invaders were quantified by the maximal OD over 24 hours of growth. The growth rates of invaders were quantified by fitting the slopes of growth curves between the two horizontal dashed lines in the figure on logarithmic scale for biomass. There is no statistically significant correlation between invasion probability of invaders with their carrying capacities and growth rates, under both high nutrient and low nutrient. The phylogeny of invader 1 to invader 9 are: *Flectobacillus*, *Pseudomonas*, *Pedobacter*, *Pseudomonas*, *Pantoea*, *Bacillus*, *Enterobacterales*, *Pantoea*, *Chryseobacterium*.

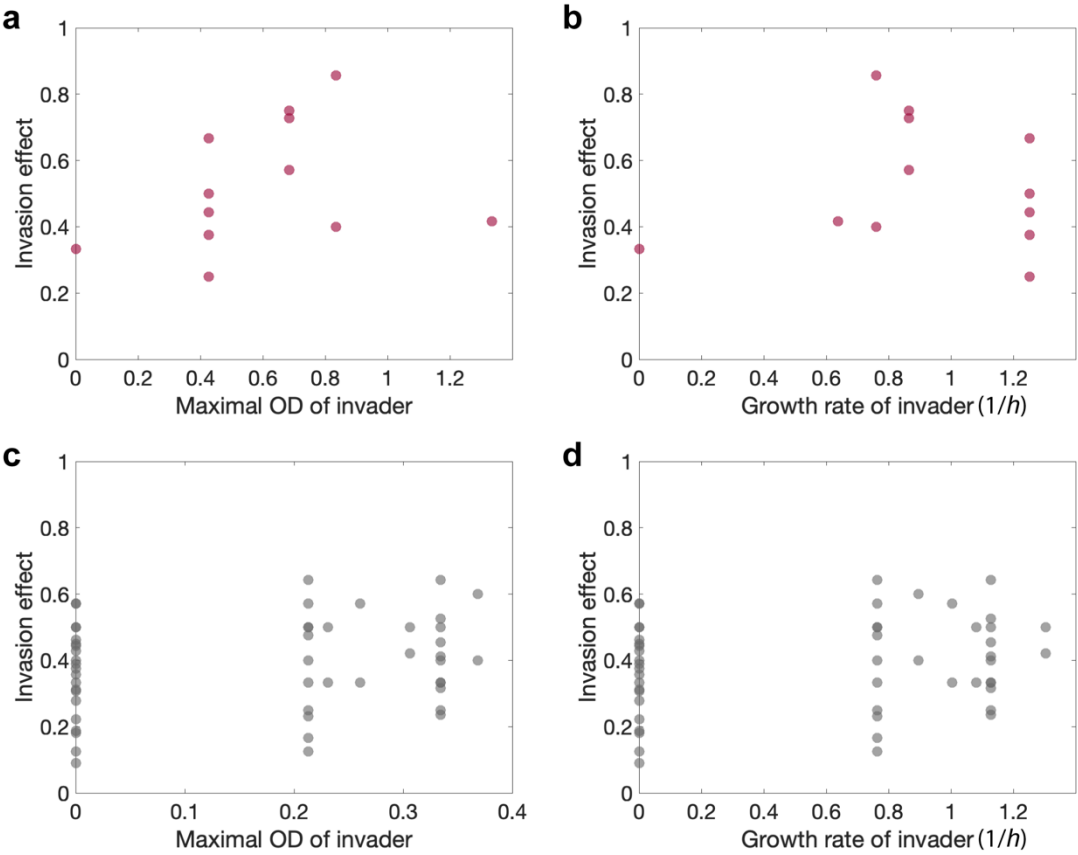

**Supplementary Fig. 20. There is no statistically significant correlation between invasion effect and invader properties.** Under high nutrient, invasion effect does not show statistically significant correlation with carrying capacity (a) and growth rate (b). Under low nutrient, invasion effect does not show statistically significant correlation with carrying capacity (c) and growth rate (d).

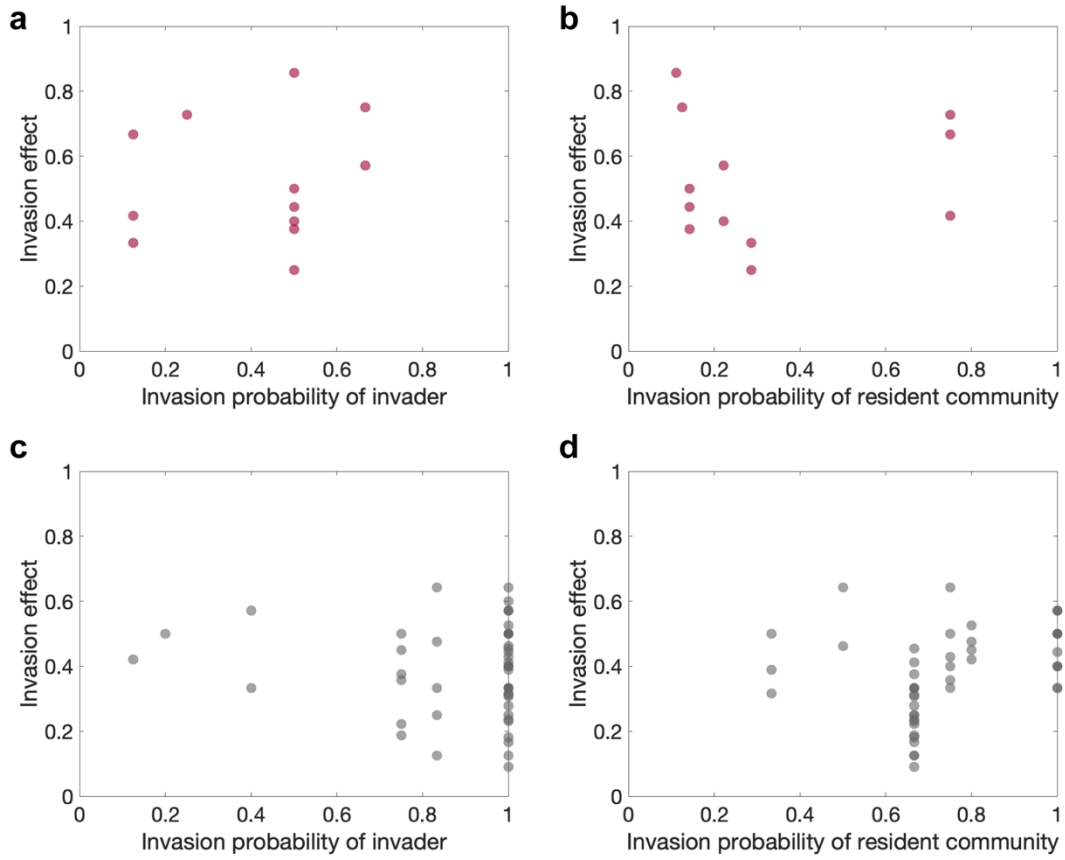

**Supplementary Fig. 21. There is no statistically significant correlation between invasion effect and invasion probability.** Under high nutrient, invasion effect does not show statistically significant correlation with invasion probability of invaders (a) and invasion probability of resident communities (b). Under low nutrient, invasion effect does not show statistically significant correlation with invasion probability of invaders (c) and invasion probability of resident communities (d).
